# Leaf hydraulic maze; Differential effect of ABA on bundle-sheath, palisade, and spongy mesophyll hydraulic conductance

**DOI:** 10.1101/2022.10.03.510099

**Authors:** Adi Yaaran, Eyal Erez, Carl Procko, Menachem Moshelion

## Abstract

Leaf hydraulic conductance (K_leaf_) facilitates the movement of water for transpiration, enabling continual CO_2_ uptake while the plant maintains its water status. We hypothesized that bundle-sheath and mesophyll cells play key roles in regulating the radial flow of water out of the xylem under optimal and stress conditions. To examine that hypothesis, we generated transgenic *Arabidopsis* plants that were insensitive to abscisic acid (ABA) in their bundle sheath (BSabi) or mesophyll (MCabi) cells. Both BSabi and MCabi plants showed greater K_leaf_ and transpiration under optimal conditions. Yet, the stomatal apertures, stomatal indices and vein densities of the BSabi plants were similar to those of WT plants. MCabi plants had larger stomatal apertures, a higher stomatal index and greater vascular diameter and biomass, relative to the WT and BSabi. In response to xylem-fed ABA, both transgenic and WT plants reduced their K_leaf_ and transpiration. However, leaf water potential was reduced only in the WT. The membrane osmotic water permeability (*P_f_*) of the WTs’ spongy mesophyll was higher than that of its palisade mesophyll. Moreover, only the spongy cells reduced their *P_f_* in response to ABA. ABA-insensitive spongy mesophyll cells had a low *P_f_*; whereas ABA-insensitive bundle-sheath cells had a higher *P_f_.* Palisade cells maintained a low *P_f_* at all ABA levels. ABA increased the symplastic water pathway, but its contribution to K_leaf_ was negligible. We suggest that the bundle sheath–spongy mesophyll pathway may control K_leaf_ to maintain steady-state conditions in the palisade cells and optimal whole-leaf water-use efficiency.

## INTRODUCTION

The movement of CO_2_ into the leaf and the loss of water from the leaf both occur through the stomata, creating a strong relationship between transpiration and productivity (de Wit, 1958; Fischer et al., 1998; Richards, 2000; Kemanian et al., 2005; Blum, 2009; Sperry et al., 2017; Sinclair, 2018; Xiong and Nadal, 2020). To support productivity (high transpiration), plants must efficiently conduct water from the soil to the evaporation sites in the leaf. Leaf hydraulic conductance (K_leaf_) is a substantial bottleneck accounting for approx. 30% (on average) and up to 98% of the total water resistance of well-watered plants (Sack and Holbrook, 2006). K_leaf_ is the ratio of water flow through the leaf (xylem to evaporation site) to the water potential gradient across the leaf (Sack and Holbrook, 2006). K_leaf_ is a dynamic trait, which responds to both abiotic (Tyree et al., 2005; Blackman et al., 2009; Shatil-Cohen et al., 2011; Pou et al., 2013; Prado and Maurel, 2013; Sade et al., 2015; Kelly et al., 2017) and biotic stress (Attia et al., 2019; Dalal et al., 2020). The structural-functional regulation of leaf hydraulics involves four main pathways: 1) hollow xylem elements; 2) the apoplast, along the cell walls of the leaf tissues; 3) the symplast, which connects cells via plasmodesmata and 4) transmembrane transport, mainly via aquaporins (AQPs). However, little is known about the relative contributions of the different pathways and the mechanisms that regulate K_leaf_.

A simplified approach divides K_leaf_ into two parts: xylem hydraulic conductance, K_x_, known as the axial path, and out-of-xylem hydraulic conductance, K_ox_, known as the radial path. Leaf vein architecture and density greatly affect K_x_ (Nardini et al., 2005a; Brodribb et al., 2007; Blackman et al., 2010; Brodribb et al., 2010; Scoffoni et al., 2011; Caringella et al., 2015). As the xylem vessels are composed mainly of dead cells, rapid impairment of K_leaf_ was previously attributed to breaks in the water-column continuum (i.e., embolism). However, in recent decades, research has revealed that living cells are involved in the regulation of K_ox_, which contributes significantly to K_leaf_ and its dynamics (Shatil-Cohen et al., 2011; Sade et al., 2014; Scoffoni and Sack, 2017; Grunwald et al., 2021).

The living vascular parenchyma, bundle-sheath cells (BSC) and mesophyll cells (MC) can potentially regulate K_ox_ according to internal and environmental signals. High K_ox_ resistance may also help leaves to avoid catastrophic xylem failure (Scoffoni et al., 2017) by dynamically regulating the resistance of the "next in line" pathway (Yaaran and Moshelion, 2016). Previously, stress-induced changes in K_ox_ were attributed to leaf shrinkage (i.e., physical alterations of the apoplastic water pathway; Scoffoni et al., 2014) and/or to changes in AQP expression and/or activity (i.e., alterations in the transmembrane water pathway). The latter has been reported in BSC (Kim and Steudle, 2009; Shatil-Cohen et al., 2011; Prado et al., 2013; Sade et al., 2015), MC (Morillon and Chrispeels, 2001) and guard cells, (Grondin et al., 2015) as well as at the whole-leaf level (Nardini et al., 2005b; Cochard et al., 2007; Pou et al., 2013; Kelly et al., 2017).

The bundle sheath is a single cell layer enwrapping the vasculature, which possesses selective barrier characteristics (Pilot et al., 2004; Leegood, 2008; Galvez-Valdivieso et al., 2009; Grunwald et al., 2021) with low levels of apoplastic transport (Shapira et al., 2009; Shatil-cohen and Moshelion, 2012). It is symplastically isolated from the xylem, which limits the transmembrane movement of water into the leaf, but is connected to neighboring MC by plasmodesmata (Ache et al., 2010). The anatomical structure of the MC correspondingly affects K_leaf_ (Brodribb et al., 2007; Scoffoni et al., 2014; Buckley, 2015). The current assumption is that the palisade MC functions as a sort of water capacitor (Zwieniecki et al., 2007); whereas the spongy MC are more dynamic in terms of their water status and play a significant role in the determination of K_ox_ (Buckley, 2015; Xiong and Nadal, 2020). In MC, water can move via all three pathways in parallel. A controlled trade-off between water pathways has been suggested as a mechanism for coping with drought stress (Morillon and Chrispeels, 2001), yet supportive evidence for this hypothesis is scarce and the evidence that does exist suggests routing toward the apoplastic pathway (Pou et al., 2013) or an increase in transmembrane pathway (Morillon and Chrispeels, 2001; Martre et al., 2002).

The drought-induced phytohormone abscisic acid (ABA) can affect transmembrane transport via the levels of expression, phosphorylation and/or degradation of AQPs (reviewed by Shivaraj et al., 2021). ABA has been shown to reduce the membrane osmotic water permeability (*P_f_*) of BSC via AQPs, to have no effect on the *P_f_* of mesophyll protoplasts (Shatil-Cohen et al., 2011) and to link the transmembrane water pathway and regulation of K_leaf_ via non-stomatal mechanisms (Pantin et al., 2013; Coupel-Ledru et al., 2017). Conversely, ABA fed through the roots was reported to increase the *P_f_* of WT mesophyll protoplasts after 24 h and that of *aba1* (mutants with impaired ABA synthesis) protoplasts after 3 h (Morillon and Chrispeels, 2001).

Focusing on the symplastic pathway, only a few works have studied the effects of ABA and abiotic stress on plasmodesmata. Those works have reported inconsistent results regarding increased (Erwee and Goodwin, 1984; Schulz, 1995) or reduced (Cui and Lee, 2016; Kitagawa et al., 2019) permeability of the plasmodesmata. We hypothesized that ABA might have a direct effect on the symplastic water pathway, in addition to its effect on the transmembrane water pathway.

In addition to its widely studied role as a stress hormone, low basal levels of ABA have recently been shown to play a major role in physiological and developmental mechanisms related to plant water status (reviewed by Yoshida et al., 2019), for example, setting the steady-state stomatal aperture (Merilo et al., 2018; Yaaran et al., 2019). However, to the best of our knowledge, the effects of basal ABA on K_leaf_ and, particularly, on K_ox_ have not yet been investigated.

The general goal of the current study was to improve our understanding of the leaf hydraulic maze, particularly along the BSC and MC continuum. We hypothesized that K_ox_ is dynamically regulated by the BSC and MC symplastic, transmembrane and apoplastic pathways in a manner that is controlled by ABA, from the low basal ABA levels observed under optimal growth conditions to the higher ABA levels that are associated with drought. To the best of our knowledge, this is the first study to investigate and compare the osmotic water permeabilities of spongy mesophyll, palisade mesophyll and BSC in the context of the regulation of leaf water balance under optimal and high-ABA conditions. To test this hypothesis, we constructed tissue-specific ABA-insensitive plants by expressing the negative dominant *abi1-1* mutant gene under a tissue-specific promoter, to generate MCabi plants (plants with ABA-insensitive MC) (Negin et al., 2019) and BSabi plants (plants with ABA-insensitive BSC), both of which maintained a WT-like stomatal response to ABA. These lines exhibited locally suppressed ABA signaling (Hoth, 2002), which may diffuse through two to seven layers of cells (Janacek et al., 2009). In this work, we address the effects of tissue-specific insensitivity to ABA from the cellular level through whole-plant transpiration rates.

## MATERIALS AND METHODS

### Plant Material and Growing Conditions

All of the *Arabidopsis thaliana* plants used in this study were of the Colombia (col) ecotype (wild type; WT). In addition to WT plants, we also used transgenic FBPase::*abi1-1* (MCabi; Negin et al., 2019) and SCR::*abi1-1* (BSabi) plants. *CalS8-1* (037603C) was generously provided by Jung-Youn Lee and *AtBG_ppap* (SAIL_115_G04) was ordered from SALK (Loughborough, UK). PCR was used to confirm the homozygosity of both of those lines. *CORI3::GUS-mCit* and *IQD22::GUS-mCit* lines were constructed by Carl Procko (Procko et al., 2022) and were crossed with MCabi#2 to generate lines with ABA-insensitive, labeled spongy or palisade MC. We labeled the bundle sheath by crossing BSabi#14 with *SCR::GFP* (Torne et al., 2021).

Arabidopsis plants were grown in 250-mL pots, with two to three plants in each pot. Those pots were filled with soil (Klasmann686, Klasmann-Deilmann; Germany) + 4 g/L Osmocote® 6M. The potted plants were kept in a growth chamber at 22°C and 70% relative humidity, with an 8-h light (09:00 – 17:00) / 16-h dark photoperiod (short day). During the day, the illumination, humidity and vapor pressure deficit (VPD) changed in a pattern resembling natural conditions, as in (Negin and Moshelion, 2017). The illumination intensity provided by LED light strips [EnerLED 24 V-5630, 24 W/m, 3000 K (50%)/6000 K (50%)] reached 150–200 μmol m^−2^ s^−1^ at the plant level at midday.

### The FBPase::*abi1-1* and SCR::*abi1-1* Constructs and Plant Transformation

For construct assembly, the MultiSite Gateway Three-Fragment Vector Construction Kit (Invitrogen; Ghent, Belgium) was used according to the manufacturer’s instructions. The construction of the MCabi plants with ABA-insensitive mesophyll is described in Negin et al. (2019). (In that work, the MCabi plants are referred as *fa* plants.) Briefly, the *abi1-1* gene (Koornneef et al., 1984) and the FBPase promoter (Lloyd et al., 1991) were cloned into pDONR plasmids and then inserted into the binary pB7M24GW (Invitrogen) plasmid. As MC and BSC are connected by plasmodesmata, we cannot exclude the diffusion of mutated *abi1-1* genes from MC to BSC. Therefore, we considered MCabi plants to be ABA-insensitive in their MC and BSC. The plants with ABA-insensitive BSC (BSabi plants) were constructed in the same way, using the SCR (Shatil-Cohen et al., 2011) promoter, which has been observed in the BSC associated with all veins in mature leaves (Wysocka-Diller et al., 2000). We cannot rule out the diffusion of *abi1-1* into other tissues in the BSabi plants, yet trans-activation of a hairpin by an enhancer trap (J1511) active in the leaf veins showed silencing spread only two to seven cells away from the vein (Janacek et al., 2009).

The dominance of the *abi1-1* mutation is due to its "gain of function" nature, negatively regulating ABA signaling (Gosti et al., 1999), with only 9% of the ABA-responsive genes remaining unaltered in *abi1-1* plants (Hoth, 2002). As such, it has a quantity-dependent attribute (high phosphatase activity represses the phosphorylation of SnRKs and, consequently, the initiation of ABA signal transduction). In our plants, *abi1-1* was constitutively expressed under a non-native promotor, in addition to the native ABI1 and other PP2Cs expression, suppressing ABA signaling. Therefore, we expected that its strongest effect would be seen at its expression site. Having said that, the dominance could be trait-dependent and, therefore, we always compared the *abi1-1* phenotype with the WT control.

Binary plasmids were floral-dipped (Clough and Bent, 1998). The presence of the transgene containing the G to A substitution was confirmed by sequencing and NocI digestion (described below). The study was performed on three independent lines of each construct, using homozygous t_3_ and t_4_ lines.

### Characterization of Transgenic Plants

#### Stomatal Aperture and Index

Stomatal aperture and index were measured as described by Yaaran et al. (2019). Briefly, epidermal peels were soaked in closure solutions under light (∼150 μmol m^−2^ s^−1^). After 1.5 h, ABA [(+)-cis, trans abscisic acid; Biosynth; Staad, Switzerland] was added to reach a concentration of 10 μM. DMSO at the same concentration was added to the control. All stomata were photographed under a bright-field inverted microscope (1M7100, Zeiss; Jena, Germany) on which a Hitachi HV-D30 CCD camera (Hitachi; Tokyo, Japan) was mounted. Stomatal images were analyzed to determine aperture size using the ImageJ software. Stomatal index was calculated as the number of stomata/number of epidermal cells in an area of 10 μm^2^.

#### Leaf Vein Density

Leaf vein density was measured as described by Grunwald (2021a). Briefly, leaves of the same age and size as those used in the K_leaf_ assay were immersed in 96% (v/v) ethanol solution and incubated at 50°C. The used ethanol was replaced with fresh ethanol until leaves were completely colorless and then was finally replaced with 88% (v/v) lactic acid solution (Fischer Scientific; UK) for overnight incubation at room temperature. Then, the leaves were rinsed in water and submerged in 0.5% safranin O dissolved in water (Sigma Aldrich cat. no S2255) for 1 min, followed by an overnight incubation in water to remove any excess dye. Then, the leaves were scanned in a water tray using an Epson 12000XL scanner (Seiko Epson; Japan) with a 2500-dpi resolution and two to three patches (∼1 cm^2^ each) for each leaf (9–10 leaves per line) were analyzed using WinRhizo™ software (https://regent.qc.ca/assets/winrhizo_about.html) for vein detection. Vein density was calculated by dividing the total vein length by the total leaf area scanned.

Leaf area is the projected area of a fully expanded leaf (average of five leaves from each plant, six plants of each of the three independent lines of MCabi and BSabi). Leaf area was assessed using the LI-3100C Area Meter (https://www.licor.com/env/).

Leaf cross-sections for anatomical analysis of the veins and mesophyll were free-hand cut using a sharp scalpel. Then, the cross-sections were cleared using ClearSee (Kurihara et al., 2015) and stained in a 1-min incubation in Toluidine Blue O dissolved in water (TB; cat. no. T-3260, https://www.sigmaaldrich.com). They were then washed with water two or three times. Cross-sections were photographed in the same set-up as that used for the analyses of stomatal aperture and index. ImageJ software was used to analyze the images to determine mesophyll length.

#### Leaf Gas-Exchange and Hydraulic-Conductance (K_leaf_) Measurements

Gas exchange and hydraulic conductance were measured as described by Negin et al. (2019), using leaves of 6- to 8-week-old plants grown under short-day conditions. Briefly, two leaves from each plant were excised before dawn and put into tubes containing artificial xylem sap (AXS; Shatil-Cohen et al., 2011) and 0.01% DMSO (control) or 10 μM ABA. The leaves were then put into hermetically sealed transparent boxes and left in the light for a minimum of 1 h. Following that period, the lids were opened for 15 min, after which time gas exchange was measured using the LI-6400xt portable gas-exchange system equipped with the Arabidopsis chamber (LI-COR, Inc.; Lincoln, NE, USA). Chamber conditions were set to 400 ppm CO_2_, PAR of 200 µE m^−2^s^−1^, VPD of ∼1.2 KPa and flow of 200 µmol/s. In the enclosed gas-exchange chamber, there is a trade-off between the stability of humidity vs. stability of the flow. We choose to keep the flow constant at the expense of humidity stability, to maintain an undisturbed boundary layer.

Immediately following the gas-exchange measurement, the leaf was transferred to a pressure chamber (ARIMAD-3000; MRC, Israel) to measure Ψ_leaf_ and determine K_leaf_ (Sade et al., 2014). K_leaf_ was calculated for each individual leaf by dividing the whole-leaf transpiration rate, E, by the leaf water potential (Martre et al., 2002; Sack and Holbrook, 2006). In our calculation, the leaf water potential gradient equals the Ψ_leaf_ measured in the pressure chamber as the leaf petiole was dipped in AXS at a water potential of 0.027 MPa. These measurements were conducted between 10:00 and 13:00 (1–4 h after the lights were turned on).

### Whole-Plant Continuous Transpiration

The whole-plant continuous transpiration rate was measured using a high-throughput telemetric, gravimetric-based phenotyping system (Plantarry 3.0 system, Plant-DiTech; Israel; Dalal, 2020) in the greenhouse of the I-CORE Center for Functional Phenotyping (http://departments.agri.huji.ac.il/plantscience/icore.phpon) as described in (Yaaran et al., 2019; Dalal et al., 2020). The output (weight) of the load cells was monitored every 3 min and analyzed using SPAC analytics (Plant-Ditech). Seedlings were transferred to 1.6-L pots (four plants per pot) and gradually exposed to semi-controlled greenhouse conditions: 24–9°C (day/night) and the natural day length and light conditions prevailing in Rehovot, Israel during December and January of 2021.

After establishment, each pot was placed on a load cell in a randomized block arrangement. Daily measurements were conducted continuously and simultaneously for all of the plants in the array, so that all of the plants were exposed to similar ambient conditions at each measurement point. The soil surface surrounding each Arabidopsis plant was covered, as well as the crack between the pot and load cell, to prevent evaporation. The transpiration rate was normalized to the total plant weight to determine E, as described in (Halperin et al., 2016; Yaaran et al., 2019).

### Protoplast Isolation and Measurement of the Membrane Osmotic Water Permeability Coefficient (*P_f_*)

Protoplast isolation and measurements of the osmotic water permeability coefficient (*P_f_*) were performed as described by Shatil-Cohen et al., (2014). In brief, protoplasts were isolated from 6- to 8-week-old plants according to the gentle and rapid (<30 min) method using an isotonic solution in the extraction process (pH 5.7, 600 mOsmol). For the ABA treatment, protoplasts were incubated in 1 μM ABA for 1–4 h. We observed the protoplasts swelling in response to the hypo-osmotic challenge (of 0.37 MPa) generated by changing the bath solutions from isotonic (600 mOsmol) to hypotonic (450 mOsmol). Palisade and spongy mesophyll protoplasts were identified by *mCit* fluorescence. Protoplasts were recorded using an inverted epifluorescent microscope (Nikon Eclipse TS100) with a 20x/NA 0.40 objective, a CCD 12-bit camera Manta G-235B (https://www.alliedvision.com) and image-processing software (AcquireControl® v5.0.0; https://www.alliedvision.com). *P_f_* was determined based on the rate at which cell volume increased for 60 s starting from the hypo-osmotic challenge, using a numerical approach in an offline curve-fitting procedure of the PfFit program.

### Relative Symplastic Permeability

#### Drop-and-See (DANS) Assays

Drop-and-see (DANS) assays were performed as described by Cui and Lee (2016) with minimal modifications. In brief, two 1-μL droplets of 2 mM 5(6)-carboxyfluorescein diacetate (CFDA; cat. #19582 https://www.caymanchem.com) were loaded on the center of the adaxial surface of a mature leaf from a 7- to 8-week-old plant. This was followed by inverted epifluorescent microscope imaging (described above for the *P_f_* measurements) of the abaxial surface of the leaf at 10 min after loading. The non-fluorescent ester CFDA can cross cell membranes passively in its electrically neutral or near-neutral form. Once inside the cells, it is subject to cleavage by esterases to form a polar fluorescent compound, CF. The extent of cell-to-cell dye movement is expected to represent the permeability of the plasmodesmata and was evaluated using an Intensity-Distance chart that was analyzed using ImageJ. A magnification of 10x was used, despite the fact that the size of some of the MCabi samples exceeded the imaging border (maximum of 388% compared to WT under control conditions), in order to capture the extent of all lines’ cell-to-cell fluorescence under the same imaging conditions. The average of the two droplets was considered the leaf’s DANS measurement.

Both attached and detached leaf measurements were performed between 10:00 and 14:00. The detached-leaf experimental set-up was like the set-up used for the measurements of hydraulic conductivity (K_leaf_).

### Statistical Analysis

The data were analyzed using Student’s *t*-test for comparisons of two means and Tukey’s HSD test for comparisons of more than two means. When two variables were examined (e.g., line and ABA treatment), the interaction between those factors was evaluated using a two-way ANOVA, followed by Tukey’s HSD test. The model effects and different *p*-values attained from the two-way ANOVAs are presented in Supplemental Table S1. All analyses were done with JMP software (SAS; Cary, NC, USA).

## RESULTS

### Characterization of the BSabi and MCabi Phenotypes

To better understand the effects of ABA on leaf hydraulics, we generated tissue-specific ABA-insensitive lines. We expressed the dominant *abi1-1* mutation under the FBPase promoter (for expression in green tissues, but not in guard cells), to create plants that had suppressed ABA signaling in their MC and BSC (i.e., MCabi plants), or under the SCR promoter (for expression in BSC, but not in MC), to create plants with suppressed ABA signaling in their BSC (i.e., BSabi plants; see Materials and Methods). *abi1-1* may diffuse through two to seven layers of cells (Janacek et al., 2009), but its strongest effect is expected at the site at which it is expressed. Three independent lines were selected for each mutation (MCabi: 1, 2 and 7; BSabi: 12, 14 and 99).

The MCabi plants were broadly characterized and validated (Negin et al., 2019) and the same validation was performed for the BSabi plants (Supplemental Figure 1). MCabi and BSabi lines exhibited different phenotypes, illustrating the vital implications of expression location. MCabi’s leaves were larger and thicker than those of the WT, while BSabi had smaller leaves (Figure 1A, F). BSabi maintained the same proportion of palisade to spongy MC as the WT; whereas MCabi had more space occupied by palisade MC (Figure 1F). Despite the changes in leaf size, the vein densities of BSabi and MCabi remained the same as that of the WT (Figure 1C), with BSabi differing from the WT in its narrower veins, but only MCabi differing from WT in its increased xylem to total vein ratio (Figure 1G). Moreover, the steady-state stomatal aperture (Figure 1D) and stomatal index (Figure E) of MCabi were greater than those of the WT and BSabi. Both MCabi and BSabi plants had a WT-like stomatal response to ABA (Negin et al., 2019; Supplemental Figure 1C) and their average foliar ABA levels were similar to those of the WT (Supplemental Figure 2A; one of the three BSabi lines had ABA levels that were higher than those of the WT).

**Figure 1.**
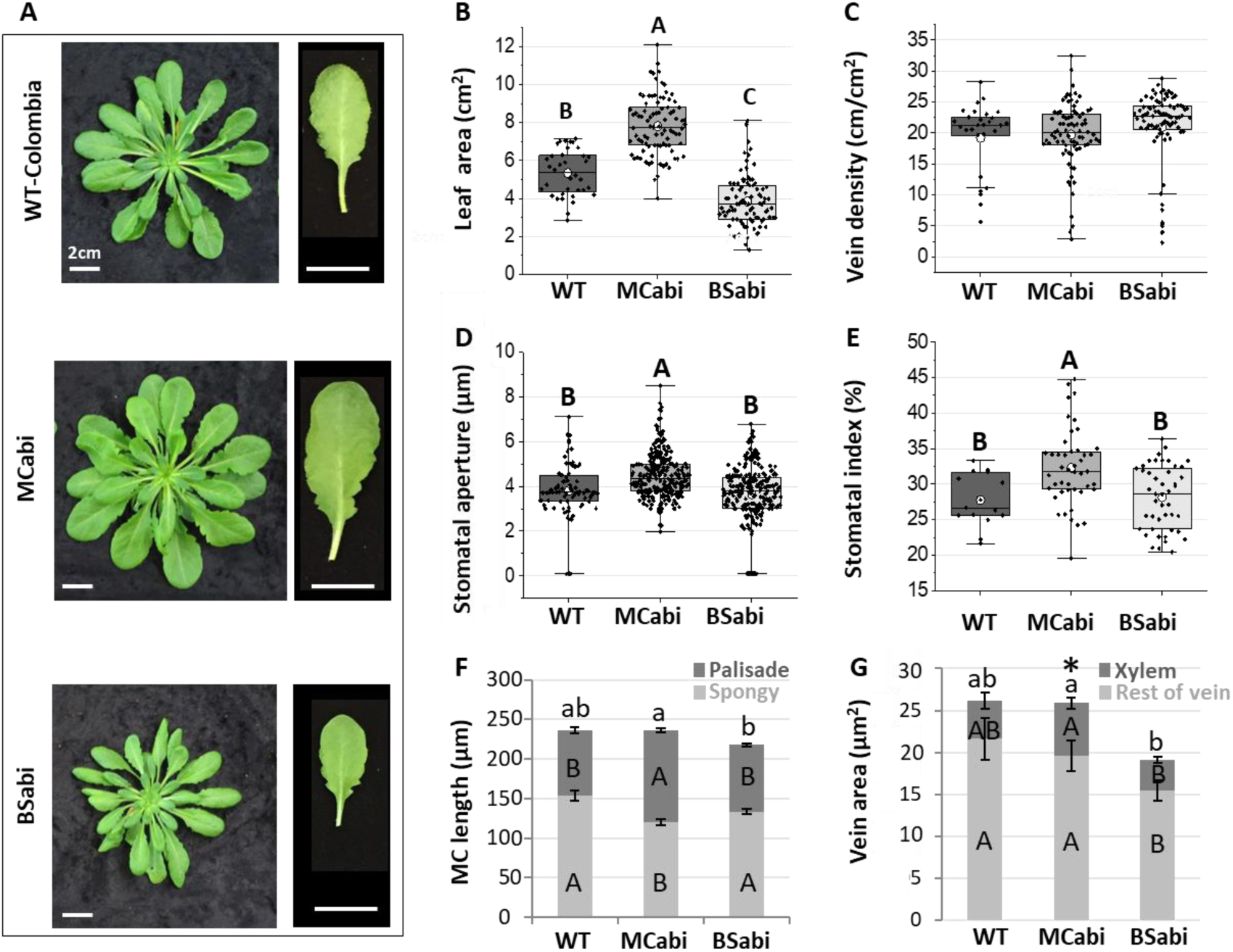
MCabi and BSabi phenotypes. Morphological characteristics of 7- to 8-week-old MCabi and BSabi plants grown under short-day conditions, including (A) representative images of the rosettes and fully expanded leaves of each line (bar = 2 cm), (B) leaf area, (C) vein density, (D) stomatal aperture, (E) stomatal index and (F) total leaf thickness divided by the widths occupied by palisade (dark gray) and spongy (light gray) mesophyll. Capital letters refers to significant differences between each mesophyll layer and lower-case letters refer to significant differences in total leaf thickness (Tukey’s test, *P* < 0.05). (G) Vein area divided into the area occupied by the xylem (dark gray) and the rest of the vein area (light gray). An asterisk indicates a significantly higher ratio of xylem to total vein area, capital letters refer to significant differences between the xylem and the rest of the vein area and lower-case letters refer to significant differences in total vein area (Tukey’s test, *P* < 0.05). (B–E) The box height shows 25–75% of the data range, the symbols indicate the individual data values, the solid line indicates the median, ᴏ indicates the mean and the whiskers delimit ±3 times the interquartile range. (F–G) Data are means ± SE. (B) *n* = 36 for WT and 90 for MCabi and BSabi, 5 leaves from 6 plants for each transgenic line. (C) *n* = 10 leaves for WT and 28–30 for MCabi and BSabi, 9–10 leaves from each transgenic line, an average of 2–3 patches of ∼1 cm^2^ per leaf. (D) *n* = 80 stomata for WT and >245 for MCabi and BSabi, ∼ 80 stomata from each transgenic line, 20 stomata from 4 leaves for each. (E) *n* = 80 stomata for WT and >245 for MCabi and BSabi, ∼ 80 stomata from each transgenic line, 20 stomata from 4 different leaves for each. (F, G) *n* = 5 leaves for WT and 15 for MCabi and BSabi, 5 leaves from each transgenic line, an average of 3 cross-sections per leaf.

### Gas Exchange and Leaf Hydraulics

To assess the relative contributions of the MC and BSC to the overall K_leaf_, we performed gas-exchange measurements and then measured Ѱ_leaf_ using a pressure chamber. The experiment was first done for each transgene separately (Supplemental Figure 3). All three independent lines were found to be similar to one another and the experiment was then repeated with MCabi and BSabi together, so that they could be compared at the same time under the same conditions (Figure 2). Under optimal conditions, BSabi and MCabi had similar levels of transpiration (E), which were higher than that of the WT. Yet, despite that higher E, both types of transgenic plants maintained Ѱ_leaf_ that was similar to that of the WT, resulting in higher K_leaf_ than the WT (Figure 2A–C). BSabi and MCabi exhibited higher stomatal conductance (g_s_) accompanied by only slightly higher net carbon assimilation rate (A_N_), resulting in lower intrinsic water-use efficiency (iWUE; Figure 2D–F). Following ABA treatment, all lines exhibited reduced E, g_s_ and, surprisingly, K_leaf_ (Figure 2A, C). MCabi and BSabi maintained an unaltered stomatal ABA response (Supplemental Figure 1C; Negin 2019) and, therefore, reduced their g_s_ and E in response to ABA (Figure 2A, D). However, while ABA reduced the Ѱ_leaf_ of the WT, the Ѱ_leaf_ levels of MCabi and BSabi remained unchanged (Figure 2B). Consequently, the K_leaf_ reduction observed in MCabi and BSabi was mainly related to their reduced transpiration (Figure 2A–C). Overall, this resulted in higher iWUE for the WT, as compared to the two sets of transgenic lines, even following ABA treatment (Figure 2F).

**Figure 2.**
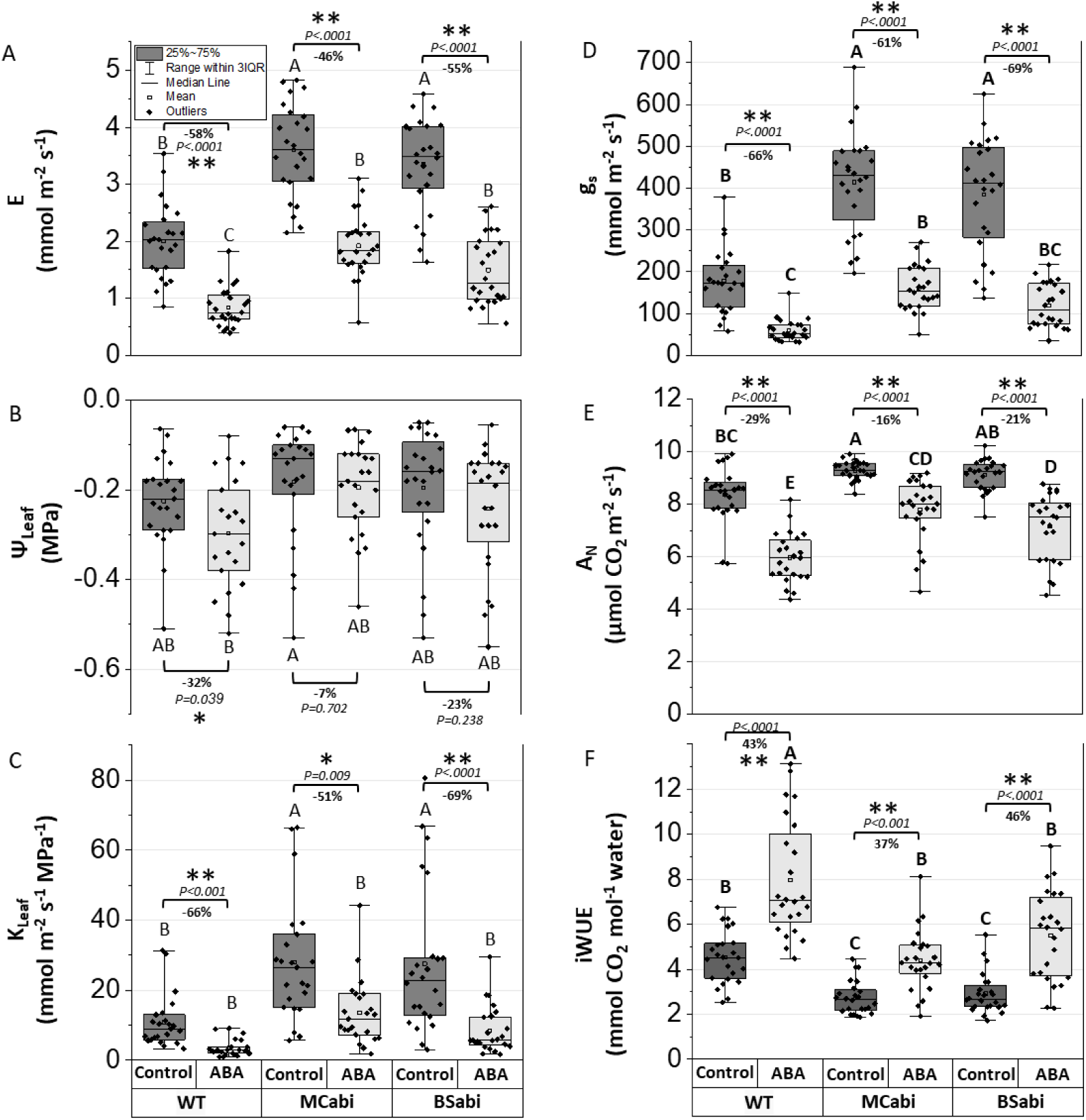
Gas exchange of WT, MCabi and BSabi leaves under non-stressed conditions in the presence of xylem-fed ABA. Leaves of 6- to 8 week-old plants were cut before dawn and petiole-fed AXS (dark gray) or AXS + 10 μM ABA (light gray). After 1–4 h, the (A) whole-leaf transpiration rate, E, and (B) leaf water potential, Ѱleaf, were measured, enabling the calculation of (C) leaf hydraulic conductance (Kleaf) for each individual leaf. In addition, the (D) stomatal conductance, gs, and (E) net carbon assimilation rate, AN, were measured, enabling the calculation of (F) iWUE for those same leaves. The percentage change following the ABA treatment is indicated as well. Data were analyzed using two-way ANOVA. (All model effects and interactions are presented in Supplemental Table 1.) Different letters indicate significant differences according to Tukey’s HSD test (*P* < 0.05); *p*-values of the effect of each treatment on each genotype according to *t*-tests are also indicated. The box height shows 25–75% of the data range, the symbols indicate individual data values, the solid line indicates the median, ᴏ indicates the mean and the whiskers delimit ±3 times the interquartile range of at least three independent experiments. *n =* 21 leaves for WT and 23–24 leaves for MCabi and BSabi, 7–8 leaves from each transgenic line.

Although MCabi plants had a higher stomatal index and larger stomatal apertures than the BSabi plants (Figure 1D, E), they had similar E and g_s_ (Figure 2A, D). E was measured in an enclosed gas-exchange chamber, in which it was difficult to maintain a constant VPD at high levels of E (Supplemental Figure 3). To overcome this challenge, we measured whole-plant transpiration in an open space, under natural conditions, using a gravimetric system in which all plants were simultaneously exposed to the same VPD. We found that the MCabi plants had the highest transpiration rate (Figure 3E). This trend was maintained throughout the whole growing period (40 days; Supplemental Figure 5C). When transpiration was normalized to plant weight, MCabi plants still had the highest E, with BSabi plants exhibiting intermediate level E and WT plants exhibiting the lowest E (Figure 3F). The finding that MCabi had the highest E suggests that the mesophyll has an additive effect on leaf water balance.

**Figure 3.**
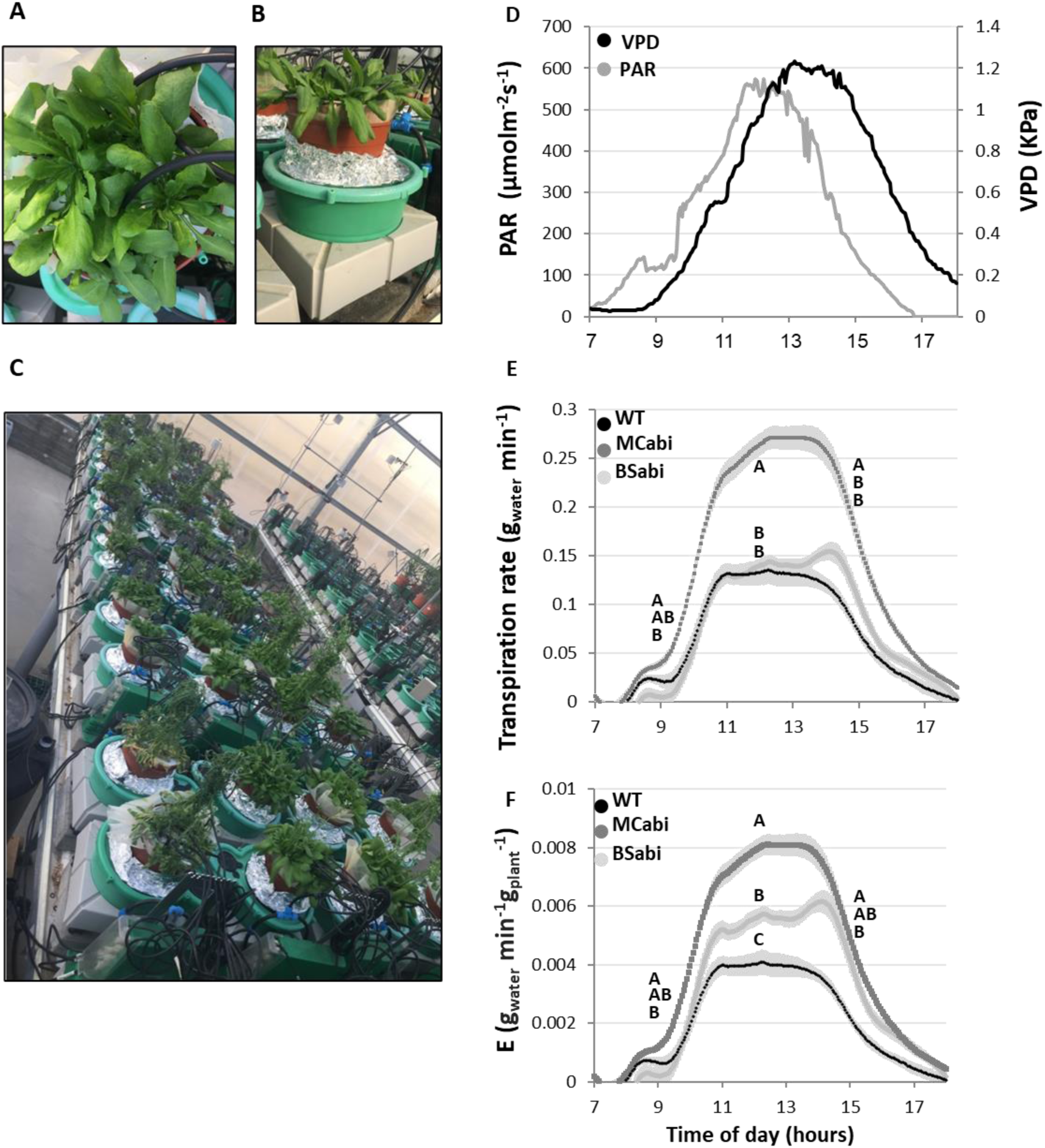
Continuous measurements of whole plants under greenhouse conditions. (A) Top view and (B) side view of pots with 8-week-old WT Arabidopsis plants, 4 plants per pot. All pots were randomized and measured simultaneously on (C) a lysimeter system located in a greenhouse under semi-controlled conditions. The plants were exposed to (D) ambient vapor pressure deficit (VPD) and PAR conditions as shown here for one representative day, the 11^th^ day after the pots were loaded onto the lysimeters. (E) Whole-plant continuous transpiration rates of WT (black), MCabi (dark gray) and BSabi (light gray). (F) Transpiration of the different lines normalized to plant weight. Data are means + SE (*n* = 5 pots for WT and 18–19 for MCabi and BSabi, 6–7 pots from each transgenic line, 4 plants in each pot). Letters indicate significant differences at 09:00, 12:00 and 15:00, according to Tukey’s test (*P* < 0.05).

### Differences Between the Palisade and Spongy MC: *P_f_* and ABA Response

To study the mesophyll’s contribution to E and K_leaf_, in general, and the specific contributions of palisade and spongy MC, we used reporter lines with palisade (*IQD22pro::GUS-mCit*) and lower spongy (*CORI3::GUS-mCit*) tissue-specific labeling with the fluorescent protein mCit (Procko et al., 2022) and examined the *P_f_* of each type of cell. The two reporter lines revealed largely restricted labeling; we did not find any apparent labeling on any other leaf tissue, including the bundle sheath and guard cells (Figure 4A, B). Spongy mesophyll protoplasts exhibited higher *P_f_* under control conditions (Figure 4C) and were smaller than the palisade protoplasts (Figure 4D). The *P_f_* values of both types of protoplasts showed a tailed distribution, with most of the protoplasts having low *P_f_* values. Yet, even when we considered only 90% of the population, to exclude the “tails” of the populations, the *P_f_* of the spongy mesophyll protoplasts remained almost double that of the palisade protoplasts. Moreover, more spongy mesophyll protoplasts had extreme *P_f_* values of over 60 µm sec^-1^. In response to ABA, the higher *P_f_* of the spongy mesophyll protoplasts was reduced by roughly 60% (19.2 ± 0.7 µm s^-1^ to 7.2 ± 1.8 µm s^-1^, respectively); whereas the average *P_f_* of the palisade protoplasts did not change (Figure 4E).

**Figure 4.**
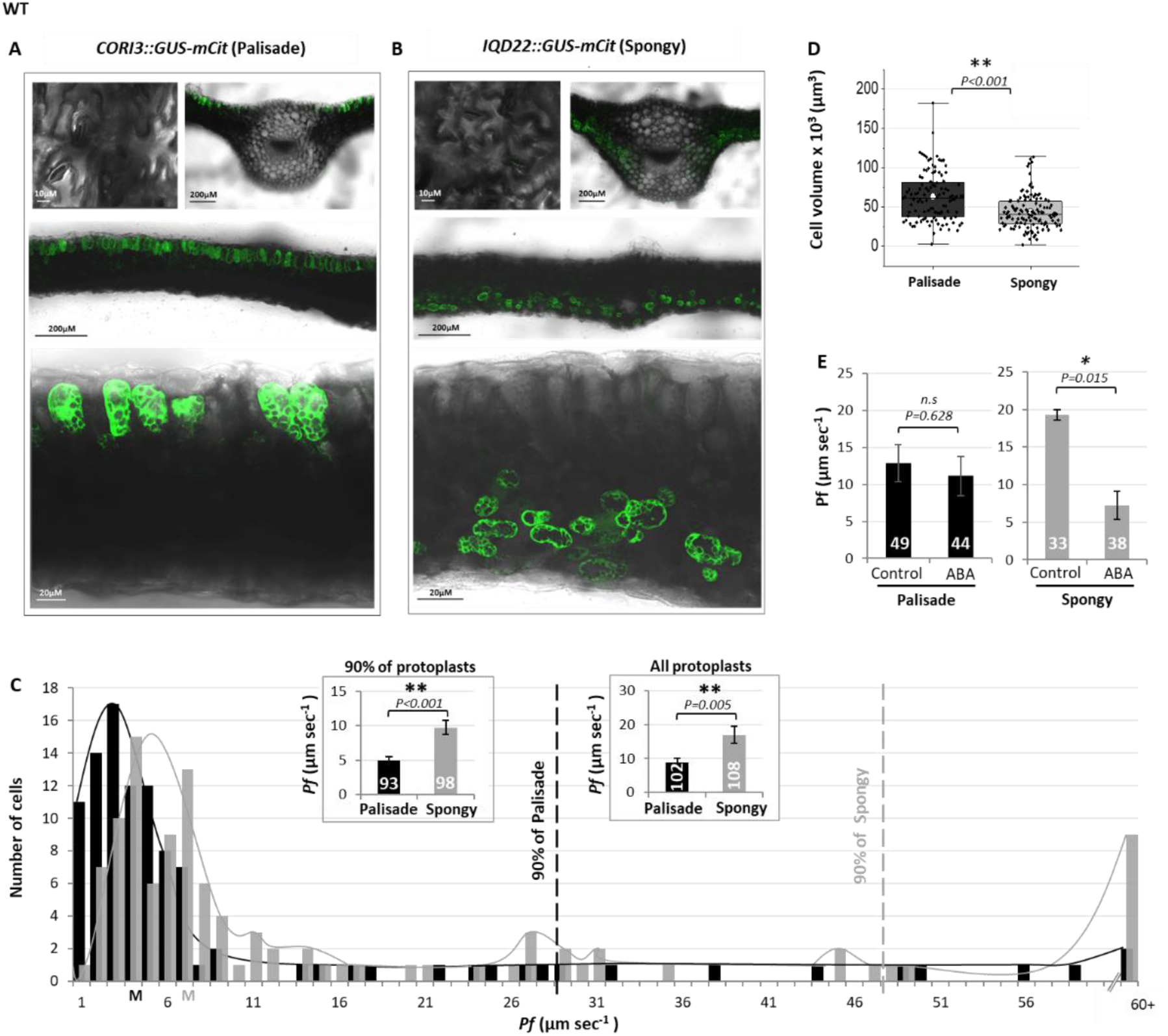
Osmotic water permeability coefficients (*Pf*) of palisade-mesophyll and spongy-mesophyll protoplasts. (A) Fluorescent images of the stomata and cross-sections of Arabidopsis leaves expressing a GUS-mCitrine reporter protein under a palisade-specific promoter (IQD22) or (B) a spongy MC-specific promoter (CORI3); bar sizes are shown in the images. (C) *Pf* distributions of WT palisade (black) and spongy (gray) MC protoplasts generated from IQD22::GUS-mCit and CORI3::GUS-mCit reporter lines, respectively. Dashed lines define the ‘tail’ of the top 10% outstanding *Pf* values of the ‘high-*Pf* ’ for both types of protoplasts (>28 µm s^-1^ for palisade and >47 µm s^-1^ for spongy). Means ± SE of 90% of the populations are also shown. M – median. (D) Volumes of palisade (gray) and spongy (black) protoplasts. The box height shows 25–75% of the data range, the symbols indicate individual data values, the solid line indicates the median, ᴏ indicates the mean and the whiskers delimit ±3 times the interquartile range. (E) *Pf* response to pre-treatment with 1 µM ABA (1–4 h). One asterisk indicates a *p*-value less than 0.05 and two asterisks indicate a *p*-value less than 0.01, according to Student’s *t*-test. The *p*-values are presented in the figure. Protoplast numbers are shown in each bar.

To examine the effect of ABA insensitivity on MC, we fluorescence-labeled palisade and spongy mesophyll protoplasts of the MCabi#2 line. Surprisingly, we found that ABA insensitivity in the spongy mesophyll reduced the *P_f_* of spongy mesophyll protoplasts, but did not affect the *P_f_* of palisade protoplasts (Figure 5A). Moreover, MCabi spongy protoplasts maintained their low *P_f_* at increasing concentrations of ABA and remained insensitive to ABA even when they were exposed to 50 μM ABA (Figure 5C).

**Figure. 5.**
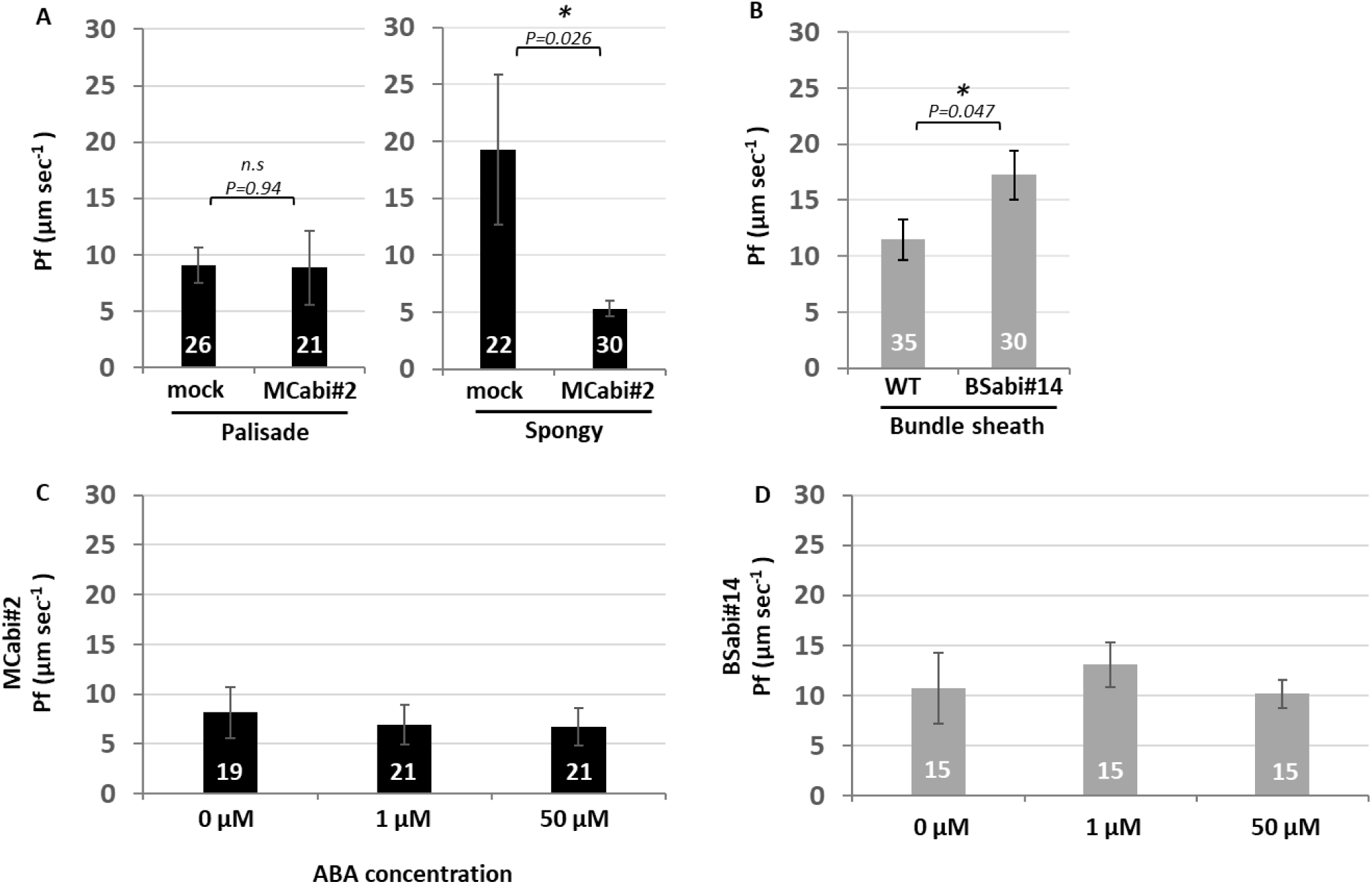
Osmotic water permeability coefficients (*Pf*) of MCabi and BSabi protoplasts in response to increasing ABA levels. (A) *Pf* levels of palisade and spongy mesophyll protoplasts of WT and MCabi#2 (ABA-insensitive) carrying the palisade (*IQD22::GUS-mCit*) and spongy (*CORI3::GUS-mCit*) markers under non-stressed conditions.. (B) *Pf* levels of bundle-sheath protoplasts of WT and BSabi#14 (ABA-insensitive) carrying the BSC marker SCR::GFP under non-stressed conditions. (C) *Pf* responses of palisade mesophyll of MCabi#2 and (D) of the bundle sheath of BSabi#14 to pre-treatment with increasing levels of ABA: 0, 1 and 50 µM ABA (1–4 h). Data are means ± SE from at least 4 independent experiments. For A and B, an asterisk indicates a *p*-value of less than 0.05, according to Student’s *t*-test. The *p*-values are presented in the figure. For C and D, there were no significant differences between the ABA concentrations, according to Tukey’s HSD test. Protoplast numbers are presented inside each bar.

We also used fluorescence-labeled ABA-insensitive BSCs of the BSabi#14 line to investigate the effect of ABA insensitivity in the bundle sheath and found that insensitivity increased the *P_f_* of the bundle-sheath protoplasts (Figure 5B). BSabi#14 bundle-sheath protoplasts also preserved their ABA insensitivity, and their *P_f_* did not change in response to increasing ABA concentration (Figure 5D), with average *P_f_* values lower than those measured in the previous experiment (Figure 5B).

We conducted another experiment with the same lines under the same conditions to determine whether this difference represents natural or experimental variation. The results of that work confirmed that WT bundle-sheath *P_f_* remained lower than the BSabi#14 bundle-sheath *P_f_*, as shown in Figure 5B (WT 8.6±2, BSabi#14 16.9 ± 6.4). Taken together, both basal and higher levels of ABA appear to affect the movement of water across spongy MC and BS membranes, affecting the transmembrane water pathway. Next, we were interested in exploring the effects of ABA on the movement of water through the symplastic pathway.

### Effects of ABA on the Symplastic Water Pathway

To evaluate the relative contribution of the symplastic water pathway to the total K_leaf_, we measured the K_leaf_ of known mutants with severely altered plasmodesmatal permeability: *CalS 8-1*, whose plasmodesmatal permeability is ∼35% higher than that of the WT (Cui and Lee, 2016) and *AtBG_ppap*, whose plasmodesmatal permeability is ∼45% lower than that of the WT (Zavaliev et al., 2013). The K_leaf_ values of both mutants were similar to that of the WT (Figure 6A) and their g_s_, E and Ψ_leaf_ were similar as well (Supplemental Figure 6). Next, we used the DANS assay (Cui and Lee 2016) to measure the relative plasmodesmatal permeability of MCabi and BSabi under control conditions and in response to ABA treatment. In principle, the cell-to-cell spread of a fluorescent dye can be used to measure the permeability of the plasmodesmata (this spread can be seen in the top of Figure 6B, marked by yellow circles. For more detailed information, see the Material and Methods). Under control conditions, the relative plasmodesmatal permeability of the MCabi plants was highest, that of the BSabi was intermediate and the WT had the lowest relative plasmodesmatal permeability (Figure 6B). This ranking corresponds to the ranking of those lines’ E under greenhouse conditions (Figure 3F).

**Figure 6.**
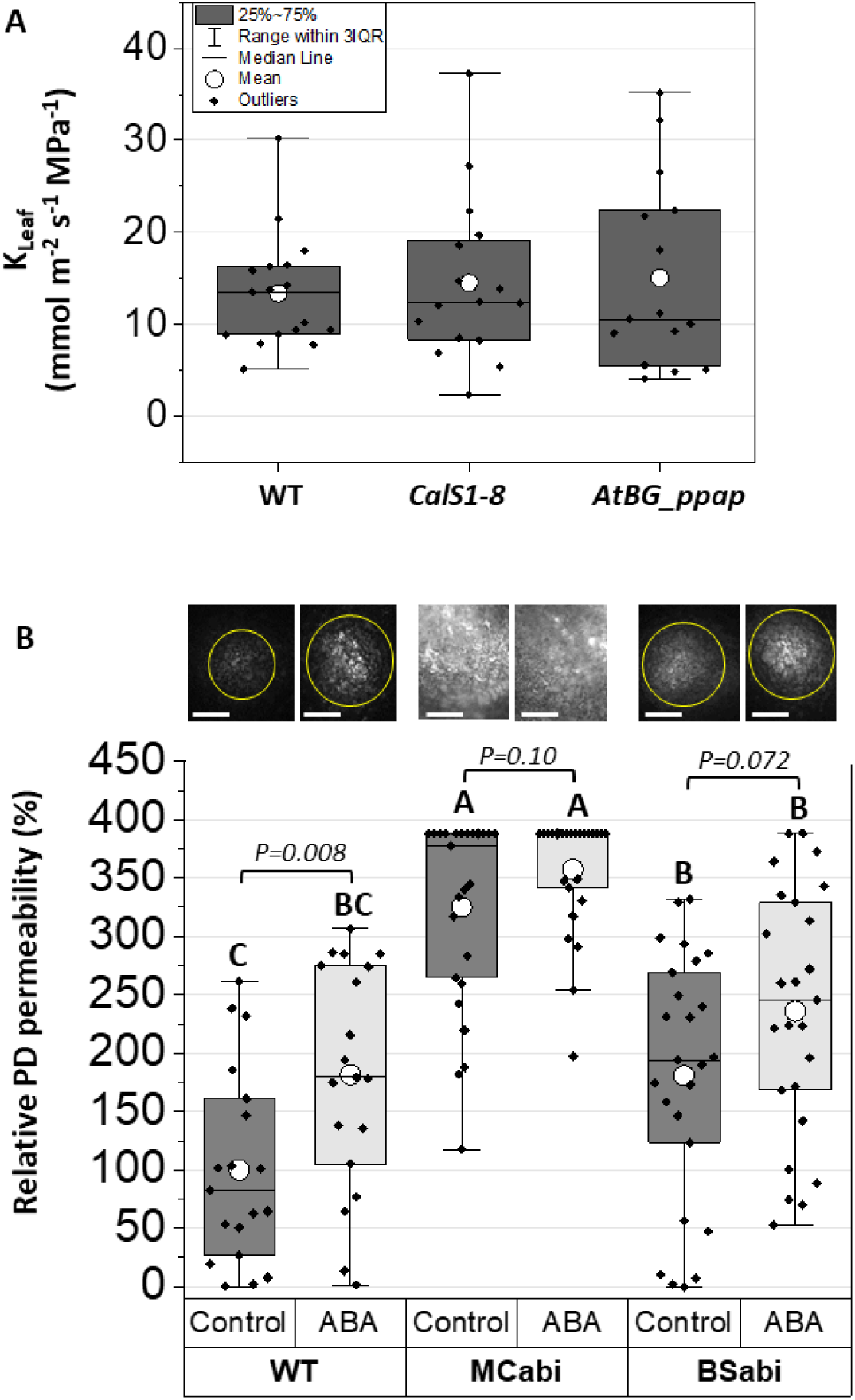
Effects of ABA on the symplastic and apoplastic pathways. (A) Kleaf of mutants with impaired plasmodesmatal (PD) permeability. (B) DANS assays revealed the cell-to-cell spread of fluorescence in WT, MCabi and BSabi plants that were petiole-fed AXS (dark gray) or AXS +10 μM ABA (light gray). Top of B: Yellow circles show the cell-to-cell spread of the fluorescent dye (bar = 20 μm). The close-up images show that the dye was trapped within the cells (bar = 5 μm). The box height shows 25–75% of the data range, the symbols indicate the individual data values, the solid line indicates the median, ᴏ indicates the mean and the whiskers delimit ±3 times the interquartile range. Different letters indicate significant differences between genotypes within a treatment (capital letters for control and lower-case letters for ABA treatment) according to Tukey’s test. An asterisk indicates a significant difference between treatments within a genotype according to Student’s *t*-test (*P* < 0.01). A: *n* = 15–17 leaves; B: *n* = 19 for WT and 24–25 for MCabi and BSabi, 8–9 leaves from each transgenic line.

Following ABA treatment, only WT leaves exhibited significantly increased relative plasmodesmatal permeability (see Figure 5B and Supplemental Figure 8A for detailed information for each line). Changes in plasmodesmatal permeability can be attributed to the accumulation of callose (a β-1,3 glucan) in the plasmodesmata (Amsbury, 2017). Nevertheless, there were no detected changes in the callose levels in either of the lines in any of the treatments in our experiment (Supplemental Figure 8B). Validation of proper callose staining was done on leaves that had been pre-treated with flagellin, whose reduced relative plasmodesmatal permeability was accompanied by increased callose deposition, as expected (Supplemental Figure 7).

## DISCUSSION

### BSC and MC Control K_leaf_ and Transpiration via ABA Regulation

The working hypothesis in this study was that K_ox_ is controlled by a series of living cells, including BSC and MC. Our results suggest that the *P_f_* levels of BS and spongy MC are a target of basal and higher ABA levels, while the way that they regulate K_leaf_ may not be straightforward. The increased E and K_leaf_ observed among both BSabi and MCabi under optimal conditions implies that, in the WT, E and K_leaf_ are limited by a non-stomatal mechanism, most likely involving the effect of basal ABA on physiological and developmental regulation.

Recently, it has been suggested that the basal levels of ABA expressed under non-stress conditions may limit the steady-state stomatal aperture (Merilo et al., 2018; Yaaran et al., 2019). Our finding that BSC insensitivity to ABA results in increased *P_f_* (Figure 5B) together with BSabi’s elevated K_leaf_ under control conditions (Figure 2C) suggest that ABA may play a similar role in BSC-specific regulation. We suggest that this dual restricting role (in the guard cells and in the BSC) of basal ABA serves to preserve leaf water balance by balancing leaf water influx (via the bundle sheath) and efflux (via guard cells) while increasing the *P_f_* of the spongy MC (Figures 4E and 5A), to sustain overall water flow and content between the controlled entry and exit positions. We further suggest that this role of basal ABA is augmented by its activity under stress conditions, when high ABA levels reduce both stomatal apertures and K_leaf_. It is interesting to note that similarities between the regulation mechanisms of guard cells and BSC have also been reported in the context of blue light signal-transduction pathways, which initiate stomatal opening while increasing bundle-sheath P*_f_* (Iino et al., 1985; Zait et al., 2017; Torne et al., 2021; Grunwald, 2022). Such a role for the bundle sheath also reinforces its importance as a leaf hydraulic checkpoint (Ache et al., 2010; Griffiths et al., 2013; Caringella et al., 2015).

Nevertheless, we wondered about the benefits of restricting hydraulic conductance (and possibly productivity) under optimal growth conditions. The answer to this question may lie in the reduced iWUE of MCabi and BSabi, which both exhibited a major increase in E as compared to WT [168–180% under controlled growth-chamber conditions (Figure 2A) and 68–80% under open-air greenhouse conditions (Figure 3F)], accompanied by a minor increase in the carbon assimilation rate, A_N_ (∼10%, Figure 2E). As E increases linearly with stomatal aperture, while A_N_ has a maximal saturation point, the optimized WUE falls within the non-maximal gas-exchange range (Yoo et al., 2009). Brodribb et al.’s (2005) idea that the photosynthetic system is saturated at high K_leaf_ levels further supports the potential disadvantage of very high K_leaf_. Therefore, non-maximal hydraulic conductance and E under optimal growth conditions may have a selective advantage at the evolutionary time scale, preventing excessive water loss that would provide only minimal gains in productivity.

Under ABA treatment, both BSabi and MCabi maintained high, unaltered Ѱ_leaf_ (Figure 2B), indicating that BSC and MC contribute to Ѱ_leaf_ regulation. Still, these plants exhibited strong reductions in E (46–55%), resulting in K_leaf_ reduction. This is unlike the results presented by Shatil-Cohen et al., (2011), in which ABA caused only a small reduction in E (22–24%) and failed to reduce K_leaf_. This demonstrates the dual effect of ABA (Pantin et al., 2013): a non-direct hydraulic mechanism enhancing stomatal closure via Ѱ_leaf_ regulation (Shatil-Cohen et al., 2011) along with a direct effect on stomatal closure. These results, although surprising, cannot rule out the importance of the bundle sheath as an internal barrier in the leaf, but rather suggest that the MC, especially the spongy MC, acts together with the bundle sheath in a series.

### Palisade and Spongy MC Differ in Their Hydraulic Properties

To better understand MC’s contribution to leaf hydraulics, we examined palisade and spongy MC separately. Palisade and spongy MC differ in both their structure and their hydraulic properties (Figure 4; (Canny, 2012; Buckley et al., 2015; Álvarez-Arenas et al., 2018; Muries et al., 2019). The smaller spongy MC are distant from each other, forming a tissue with a large surface area, air spaces and low cell-to-cell connectivity (Buckley et al., 2015; Borsuk et al., 2022). Based on our findings, the *P_f_* of the spongy MC under non-stressed conditions was almost twice that of the palisade MC (Figure 4C, E). The spongy mesophyll’s vast exposure to the leaf’s internal air spaces together with the high *P_f_* of its cells may explain those cells’ rapid loss of water and shrinkage in response to high VPD (Canny, 2012) or dehydration (Muries et al., 2019). This is an intriguing point, which may suggest that spongy MC act as hydraulic sensor in the following way: High *P_f_* enables the rapid loss of water content, resulting in decreased cell volume, which lessens cell-wall and plasma-membrane interactions, to induce ABA production (Bacete et al., 2022). In this way, a physical change, such as high VPD causing an increased rate of water loss from the leaf, can be translated into a physiological response of stomatal closure (Mott, 1991). Surprisingly, MCabi exhibited high K_leaf_ (Figure 2C) despite its spongy MC’s low *P_f_*, suggesting that its high K_leaf_ is mainly supported by a bundle sheath that is insensitive to ABA and apoplastic flow, as concluded from its ability to maintain high Ψ_leaf_ (Figure 5B). Nevertheless, the low *P_f_* of MCabi’s spongy MC may obstruct their hypothesized role as a hydraulic sensor (mentioned above), resulting in their higher E, g_s_, Ψ_leaf_ and K_leaf_ (Figure 2A–D). According to these results, spongy MC *P_f_* dynamics may have a regulatory role rather than directly changing K_ox_. In response to higher ABA levels, the WT spongy MC reduced its *P_f_* to avoid further water loss, while the palisade *P_f_* remained unchanged (Figure 4E), highlighting tissue-specific ABA regulation.

But why would ABA-insensitive spongy MC present a low *P_f_* similar to that of WT spongy mesophyll exposed to ABA (Figures 5A and 4E, respectively)? To answer this question, we propose that spongy MC behave according to an ABA optimization curve (for further elaboration, please see Supplemental Figure 9). Palisade MC’s lack of response to ABA supports the idea that these cells play a static hydraulic role (Zwieniecki et al., 2007), in which they serve as a water buffer to maintain leaf water balance (Nardini et al., 2010; Rockwell et al., 2014). The combination of palisade mesophyll with static hydraulic properties and the lack of any direct effect of ABA on photosynthesis (Negin et al., 2019) can be an advantage for continued photosynthesis (Tholen et al., 2012; Buckley, 2015) and position ABA as, primarily, a water-balance regulator (from BSC to guard cells) that controls the transmembrane water pathway.

### The Role of Plasmodesmata in the Symplastic Water Pathway and K_ox_

To evaluate the roles of plasmodesmata and the symplastic water pathway in K_leaf_, we used impaired plasmodesmata-permeability mutants (*cals8-1*; (Cui and Lee, 2016) and *AtBG_ppap*; (Zavaliev et al., 2013)). Both of those mutants exhibited WT-like K_leaf_ (Figure 5A), indicating that the symplastic pathway has a negligible influence on K_leaf_. Moreover, the fact that ABA increased the WT’s symplastic permeability (Figure 5B), alongside its reduction of K_leaf_, supports this idea. As high ABA levels reduce *P_f_* while increasing plasmodesmatal permeability, we suggest that the symplastic water flow may provide better cell-to-cell conductivity and turgor sharing and help to maintain a uniform water potential across the tissue. Additionally, it may act as a rapid facilitator of the stress signal by distributing the ABA among the cells.

The greater symplastic conductivity of BSabi and the even greater symplastic conductivity of MCabi under control conditions are, therefore, surprising (Figure 6B). Here, we present three possible explanations for this. The first is altered development. During sink-to-source transition, the leaves’ plasmodesmata are dramatically reduced (Roberts et al., 2001). Therefore, impaired maturation may sustain a higher number of plasmodesmata. Interestingly, *abi1-1* hybrid aspen trees fail to block their plasmodesmata as they enter dormancy (Tylewicz et al., 2018). The second possible explanation is impaired hormonal crosstalk with salicylic acid. Salicylic acid reduces the permeability of plasmodesmata (wang et al., 2013; Cui and Lee, 2016). ABI1 binds directly to salicylic acid and impairs that crosstalk (Manohar et al., 2017). The relatively low levels of salicylic acid observed in MCabi and BSabi (Supplemental Figure 2B) support the existence of such altered crosstalk, which may affect the permeability of the plasmodesmata. Finally, the *abi1-1* protein may have a signaling function. While *abi1-1* mutants lack phosphatase activity, the protein itself may have a signaling function independent of its phosphatase activity (Gosti et al., 1999). If plasmodesmatal permeability is involved in such an ABA-signaling pathway, it may be boosted in the transgenic plants due to the high abundance of *abi1-1* in the specific tissue. Although compelling, we did not pursue these leads as they go beyond the scope of this study. Further research is needed to determine ABA’s effect on the permeability of plasmodesmata.

### A Suggested Model for the Role of ABA in the Regulation of K_leaf_

Integrating our results, we suggest the following hypothetical model of radial water movement within the out-of-xylem leaf tissues (Figure 7). Under optimal conditions, water enters the leaf through the BSC, which selectively regulate transmembrane radial water flux (Leegood, 2008; Galvez-Valdivieso et al., 2009; Shapira et al., 2009; Ache et al., 2010; Shatil-Cohen et al., 2011; Prado and Maurel, 2013; Grunwald et al., 2021) based on the osmotic water permeability of their membranes (Figure 5B). Thus, the bundle-sheath barrier serves as the first control point, regulated by basal ABA (Figure 7A, Control Point 1). Then, water proceeds toward and within the MC, with most of the water flowing through the hydraulically dynamic spongy MC (Figure 4C; Buckley et al., 2015; Buckley, 2015; Xiong and Nadal, 2020). The transmembrane pathway (Cochard et al., 2007; Kim and Steudle, 2007) and/or the apoplastic pathway (Sack and Holbrook, 2006; Voicu et al., 2009; Buckley, 2015) govern K_leaf_ under optimal conditions, while the contribution of the symplastic pathway is negligible (Figures 6 and 7A, Control Point 2).

**Figure 7.**
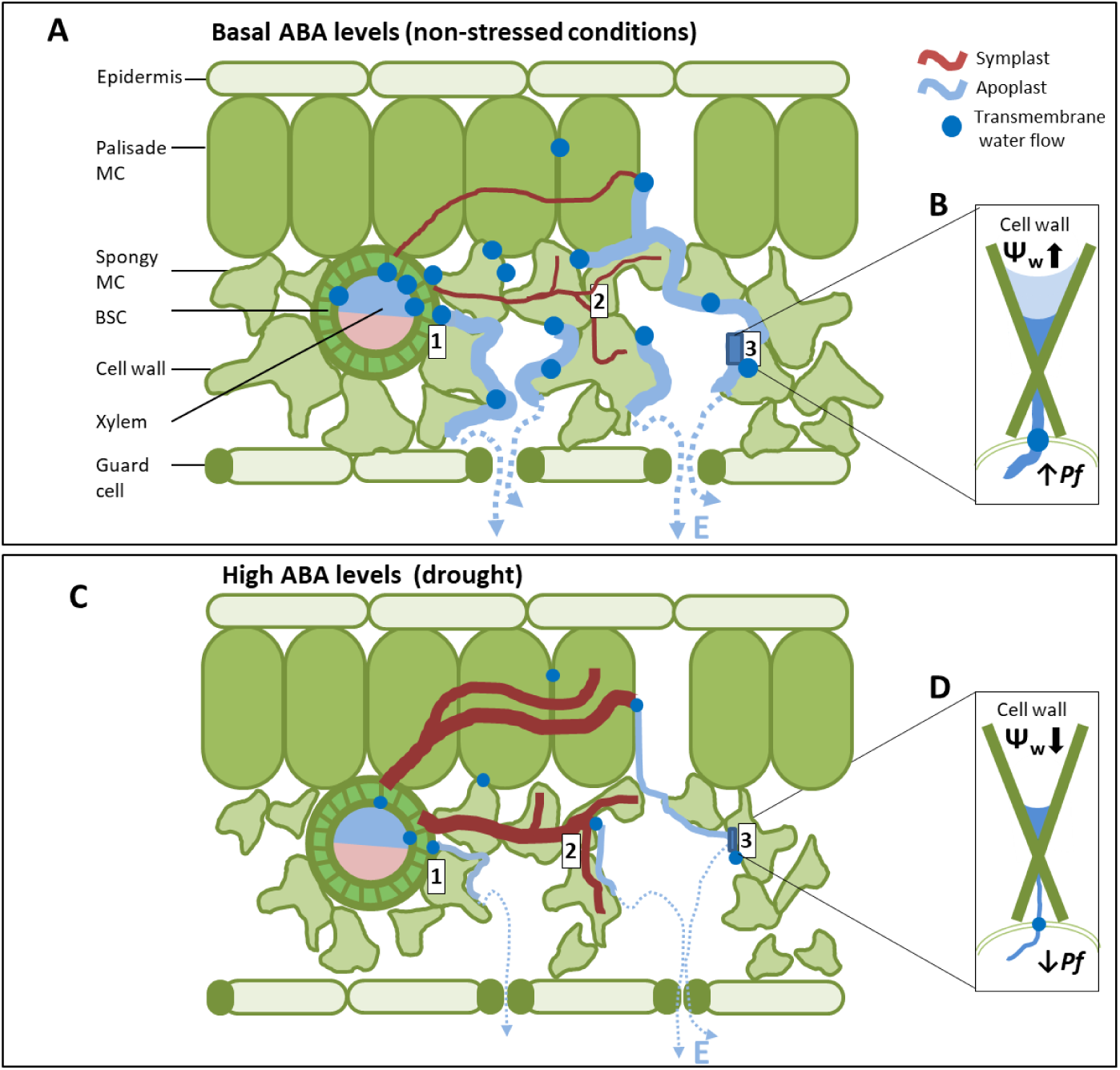
A suggested model for ABA’s regulation of Kleaf. (A) Under optimal conditions, water flows across the bundle sheath via the transmembrane water pathway, probably through aquaporins (blue dots; Control Point 1) and proceeds toward the mesophyll. In the mesophyll, water may flow in parallel pathways and move between them. The apoplastic (light blue line) and transmembrane pathways are dominant, while the contribution of the symplastic path (dark red line) is relatively small (Control Point 2). (B) The transmembrane water permeability (*Pf*) of the spongy mesophyll cells can facilitate the export of water from cells, contributing to transmembrane water flow or directing water toward the cell wall to support apoplastic water flow (Control Point 3 (and transpiration (E, dashed blue line). Note that under optimal conditions, most water flows through the spongy mesophyll cells, preserving leaf water status (e.g., high Ѱleaf and turgor). (C) High levels of ABA reduce the *Pf* of the bundle-sheath cells (blue dot, Control Point 1. Shatil-Cohen et al., 2011). In the immediate term, less water enters the leaf while the amount of water leaving through the stomata remains high, which causes the spongy mesophyll cells to shrink (Canny, 2012) due to their high initial *Pf*, amplifying the stress signal and initiating ABA production (Bactete, 2021). Higher ABA then decreases the spongy mesophyll cells’ *Pf* and (D) decreases the transmembrane water flow and apoplast wetting (Control Points 2 and 3). Diminished flow over the transmembrane and apoplastic water pathways (blue dots and light blue line, respectively) keeps water inside the cells, while ABA increases the amount of water transported via the symplastic pathway (red line; Control Point 2). The effects of all the above combined with the direct effect of ABA on stomata result in reduced Kleaf and E (dashed blue line) as the leaf’s water status deteriorates (e.g., reduced Ѱleaf and turgor).

The high *P_f_* of the spongy MC may have two advantages, allowing the spongy mesophyll to serve as an interchange point between the parallel water pathways and as a hydraulic sensor. As an interchange point between the parallel water pathways, water crossing a cell membrane may continue toward the next cell in the transmembrane pathway or remain in the apoplast, wetting the cell wall to support the apoplastic water pathway (Maurel, 1997; Steudle and Peterson, 1998; Figure 6A, Control Point 3) and transpiration (Figure 3F and Supplemental Figure 5). As a hydraulic sensor, the spongy mesophyll’s surface area combined with its high *P_f_* results in rapid water loss when there is an imbalance between water supply and demand, quickly reducing the cell’s water content and volume and offering a conceivable mechanism to capture changes in the rate of water loss from the leaf. The simultaneous limitation of K_leaf_ and stomatal aperture by basal ABA levels maintains the balanced leaf water status that is mirrored in the high Ψ_leaf_ and turgor (Figure 7A).

In the presence of a high level of ABA, xylem ABA reduces the BSC’s *P_f_* (Shatil-Cohen et al., 2011; Figure 6C, Control Point 1), which limits the movement of water into the leaf. In the immediate term, the stomatal efflux remains unchanged and the spongy MCs’ high initial *P_f_* (Figure 4C) results in their rapid water loss, shrinkage (Canny et al., 2012; Muries et al., 2019) and ABA production (McAdam and Brodribb, 2016; Sussmilch et al., 2017; Sack et al., 2018; Bacete et al., 2022; Figure 7A, B). When they sense ABA, the spongy MCs decrease their *P_f_* (Figures 4E and 6C, Control Point 2, Figure 7B), resulting in the retention of water within the whole mesophyll tissue and reducing the flow over the transmembrane water pathway, cell-wall wetting and, subsequently, the apoplastic flow (Figure 7C, D, Control Point 3). As a result of all the above and the direct effect of ABA on stomata, K_leaf_ and g_s_ decrease, while the leaf water status deteriorates (e.g., decreased turgor and Ѱ_leaf_). Simultaneously, the high ABA levels increase the symplastic permeability (Figures 6B and 7C, Control Point 2). This may facilitate the cooperation of mesophyll cells and solute transport when E levels are low (Morillon and Chrispeels, 2001) and may enhance the symplastic passage of ABA through the leaf and toward the guard cells.

This study highlights the differential effects of ABA on BSCs and spongy and palisade MCs, as well as newly reported effects of those tissues on K_leaf_. Our efforts to untangle the effects of ABA on leaf hydraulics have revealed that a series of living cells, which are differentially affected by ABA, take part in determining leaf water balance under optimal and high-ABA conditions. We also provide evidence that supports the hypothesis that ABA acts as a central water-balance coordinator and suggest that ABA levels dynamically fine-tune the movement of water into and out of leaves.

### Acknowledgments and Funding

This research was supported by the Israel Science Foundation (Grant No. 1043/20) and NIH (Grant No. 5R35GM122604). We thank Jung-Youn Lee for generously providing us with the *CalS8-1* (037603C) line. We thank Dr. Julius Ben-Ari for the LC-MS/MS analyses of ABA samples and Eduard Belausov for the confocal imaging.

### Author Contributions

Adi Yaaran designed and performed the research, analyzed the data, formulated the hypotheses and wrote the manuscript. Erez Eyal contributed to the characterization of transgenic plants and the *P_f_* measurements. Carl Procko constructed and validated the *CORI3::GUS-mCit* and *IQD22::GUS-mCit* lines. Menachem Moshelion (the corresponding author) formulated the hypotheses and wrote the manuscript together with Adi Yaaran.

