## Supplementary Figures for "Leaf hydraulic maze; Differential effect of ABA on bundle-sheath, palisade, and spongy mesophyll hydraulic conductance"

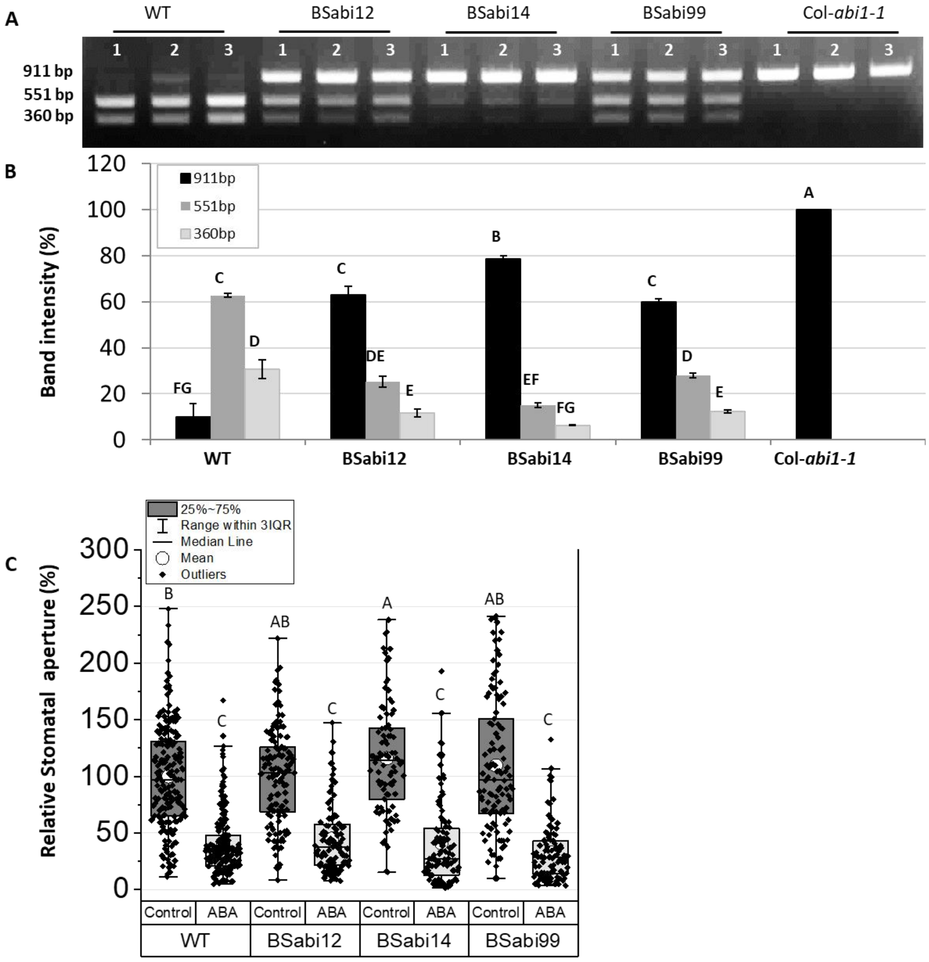


**Supplementary Figure 1.** Characterization of BSabi plants. (A) Semi-quantification of the native ABI1 gene vs. *abi1-1* mutant gene in WT, BSabi and col-*abi1-1* leaves by gel electrophoresis of the PCR-amplified ABI1 gene fragment followed by digestion by the NocI restriction enzyme. The undigested fragment (*abi1-1*) was 911 bp long and digestion (ABI1) yielded 551-bp and 360-bp fragments. (B) The intensities of the bands for each plant were quantified and expressed as a percentage. (C) Stomatal apertures of WT, BSabi12, BSabi14 and BSabi99 epidermal peels directly exposed to control (dark gray) or 10 µM ABA (light gray) solutions are presented relative to the stomatal apertures of stomata treated with control solution on each day of experiment (1.82 ± 0.06 µm), The box height shows 25–75% of the data range, the symbols indicate the individual data values, the solid line indicates the median, o indicates the mean and the whiskers delimit ±3 times the interquartile range. Different letters indicate statistically significant differences according to Tukey's test ($P < 0.05$; $n > 6$ leaves and > 90 stomata).

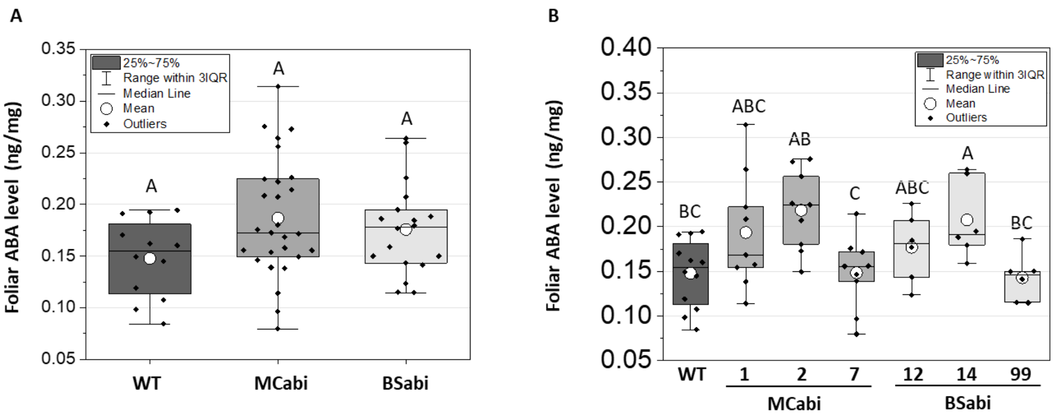


**Supplementary Figure 2.** ABA levels of MCabi and BSabi. (A) Quantification of the foliar ABA levels of well-irrigated WT, MCabi and BSabi plants, with all three independent lines of each type of transgenic plant combined and (B) with the data for each line presented separately. The box height shows 25–75% of the data range, the symbols indicate individual data values, the solid line indicates the median, o indicates the mean and the whiskers delimit ±3 times the interquartile range. Different letters indicate statistically different means (Tukey's test $P < 0.05$). A: $n = 6$ for WT; $n = 18$ for transgenic plants. B; $n = 6$.

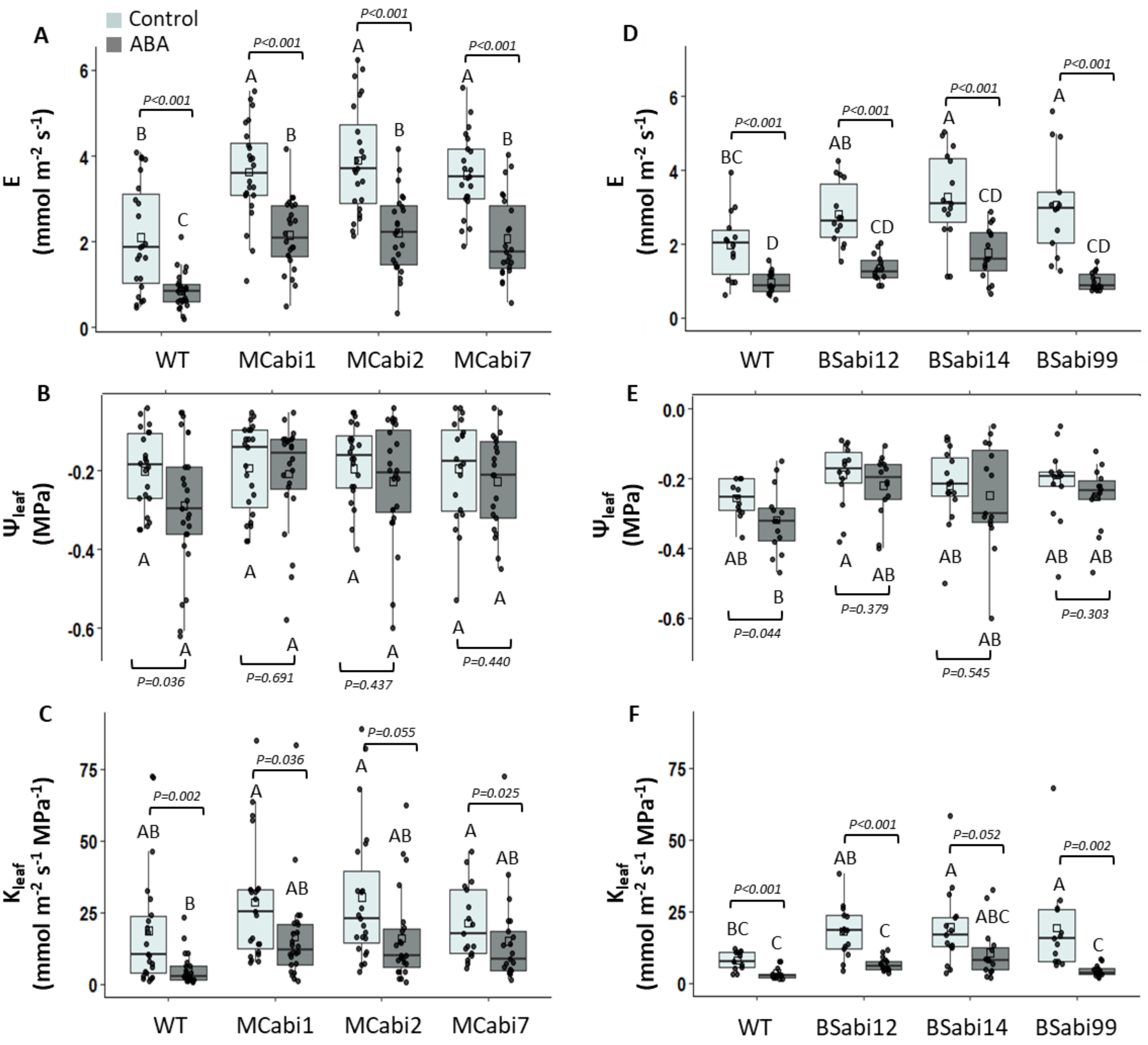


**Supplementary Figure 3.** Leaf hydraulic conductance ($K_{leaf}$) of the three independent lines of MCabi and BSabi plants. Leaves of 6- to 8-week-old plants were picked before dawn and petiole-fed AXS alone (light gray) or AXS containing 10 µM ABA (dark gray). After 1–4 h, the (A, D) whole-leaf transpiration rate, E, and (B, E) leaf water potential, $\Psi_{leaf}$, were measured, enabling the calculation of (C, F) leaf hydraulic conductance ($K_{leaf}$) for each individual leaf. Panels A–C refer to MCabi and panels D–F refer to BSabi. Data were analyzed using two-way ANOVA. (All model effects and interactions are presented in Supplemental Table 1.) Different letters indicate significant differences according to Tukey's HSD test ($P < 0.05$); *p*-values of the effects of the treatment on each genotype, according to *t*-tests, are also presented. The box height shows 25–75% of the data range, the symbols indicate individual data values, the solid line indicates the median, o indicates the mean and the whiskers delimit ±3 times the interquartile range of at least three independent experiments. A–C: $n$ = 20–25 leaves. D–F: $n$ = 13–15 leaves.

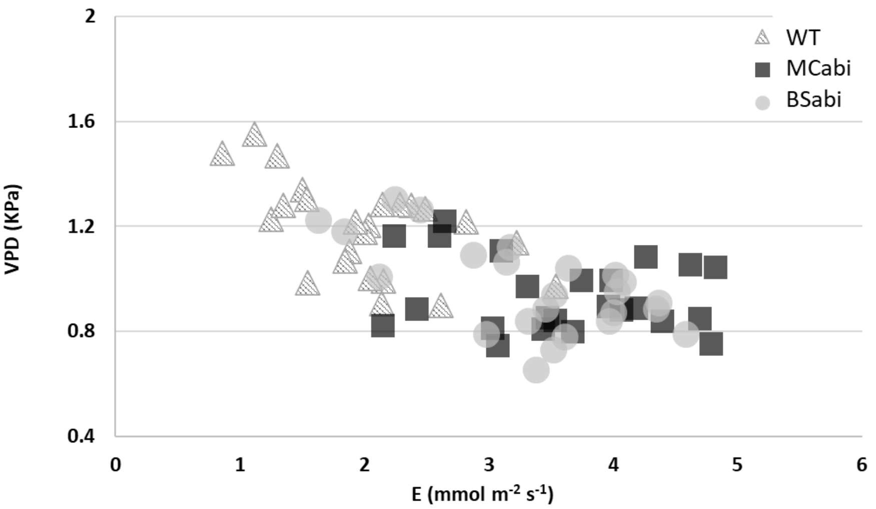


**Supplementary Figure 4.** The high transpiration of MCabi and BSabi reduced the VPD in the small gas-exchange chamber. The effect of VPD on E was examined in the gas-exchange chamber. Chamber conditions were set to 400 ppm $CO_2$, PAR of 200 µE $m^{-2} s^{-1}$, VPD of ~1.2 and a flow of 200 µmol $s^{-1}$. The flow was kept constant to avoid any diminishment of the boundary layer. $n$ = 21–24.

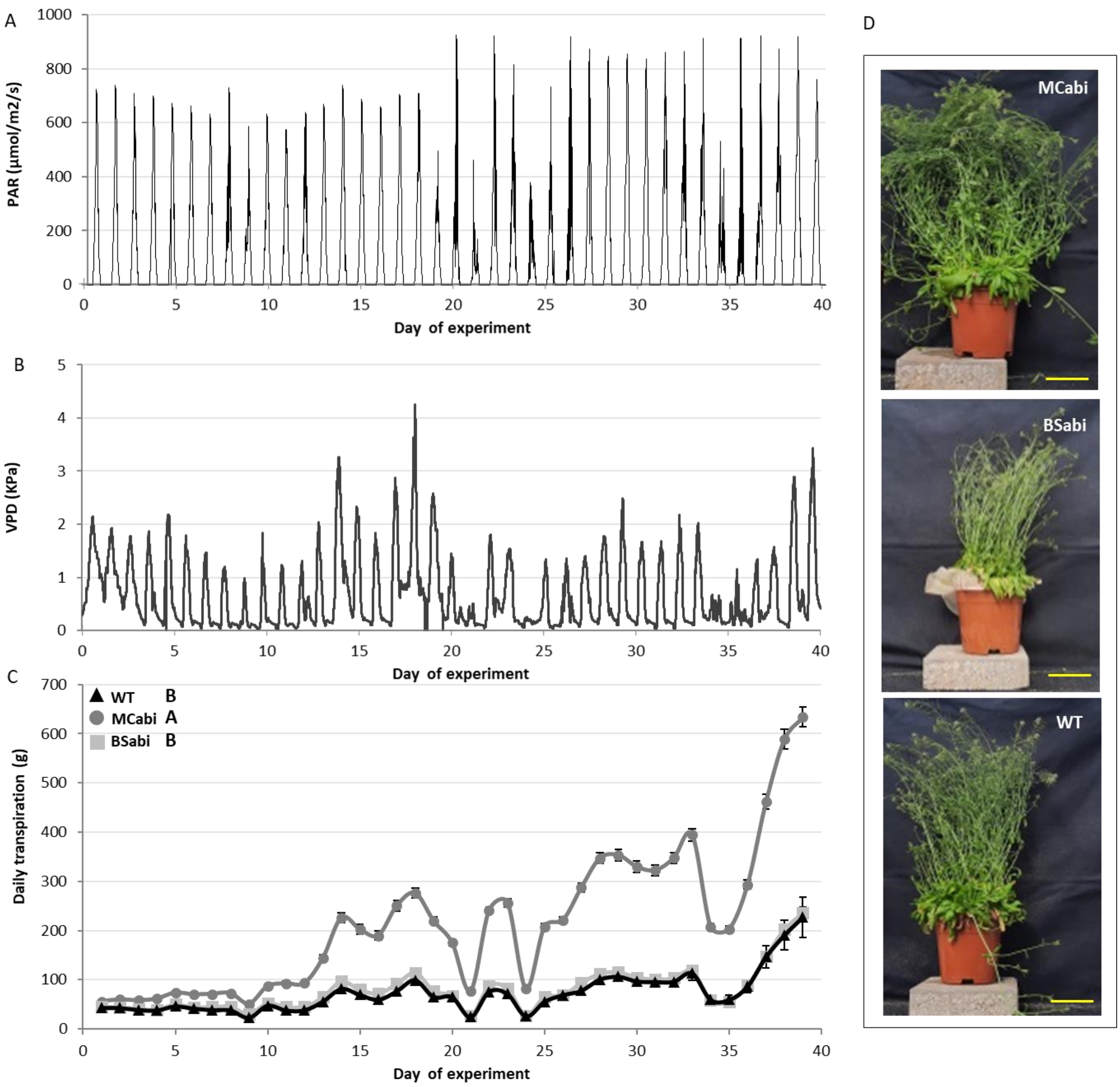


**Supplementary Figure 5.** Continuous whole-plant data collected under greenhouse conditions throughout the experiment. Natural environmental conditions of (A) PAR and (B) VPD in the semi-controlled greenhouse. (C) Whole-plant daily transpiration of WT (black triangles), MCabi (dark gray circles) and BSabi (light gray squares) plants over the whole experiment. (D) Representative picture of each genotype on the last day of the experiment; bar = 10 cm. Data are means ± SE (WT: $n$ = 5 pots; MCabi: $n$ = 19 pots; BSabi: $n$ = 18 pots; each pot held 4 plants). Different letters indicate a significant difference between genotypes according to the Tukey-Kramer test ($P$ < 0.05).

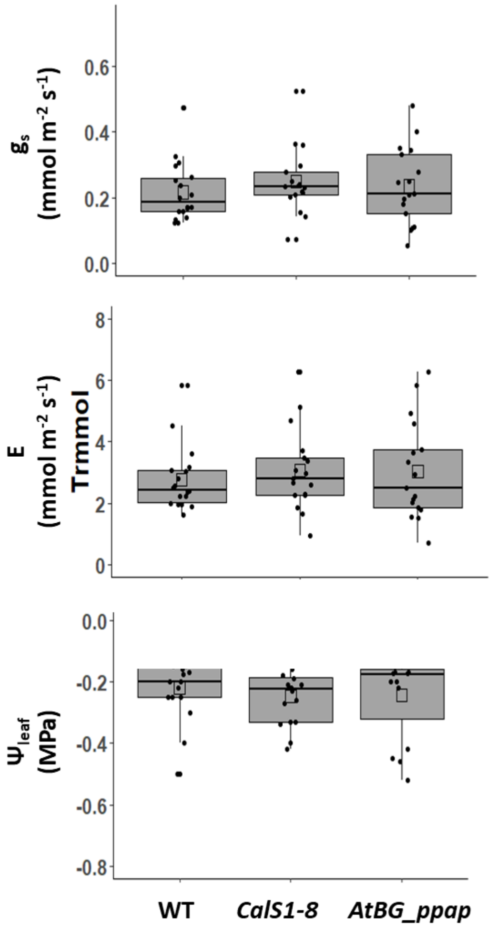


**Supplementary Figure 6.** Gas-exchange measurements of mutants with impaired plasmodesmatal permeability. *n* = 15–17 leaves. There were no significant differences between the lines, according to Tukey's HSD test.

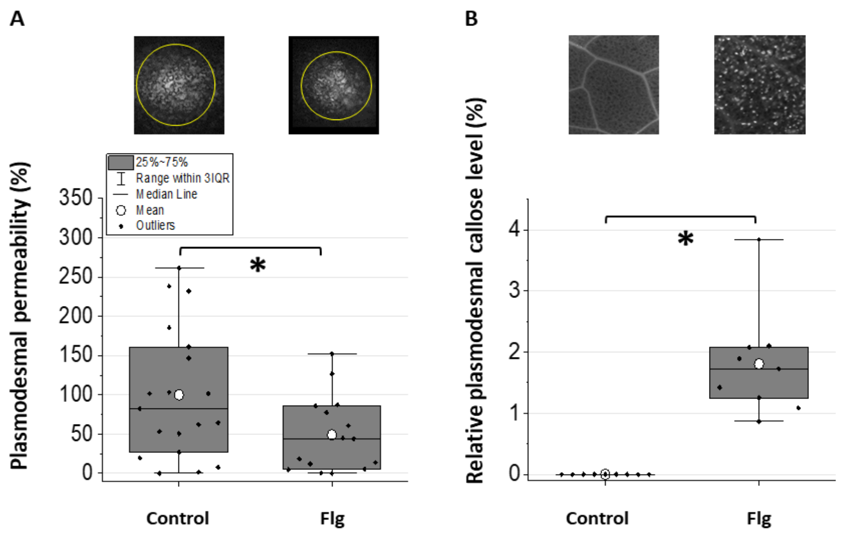


**Supplementary Figure 7.** Validation of measurements of symplastic permeability. (A) DANS assays and (B) callose staining of WT infiltrated with 20 μM flagellin 22–24 h before the measurements were taken revealed that a flagellin-dependent decrease in plasmodesmatal permeability reduced both the cell-to-cell movement of the fluorescent dye (see CFDA symplast flow) and callose deposition. Top: Yellow circles show the cell-to-cell spread of the fluorescent dye. The experimental setup is similar to that used to examine $K_{leaf}$ (Figure 2; $n = \geq 15$ leaves). The box height represents 25–75% of the data range, the symbols represent individual data values, the solid line indicates the median, o indicates the mean and the whiskers delimit ±3 times the interquartile range. An asterisk indicates a significant difference according to Student's *t*-test ($P < 0.01$).

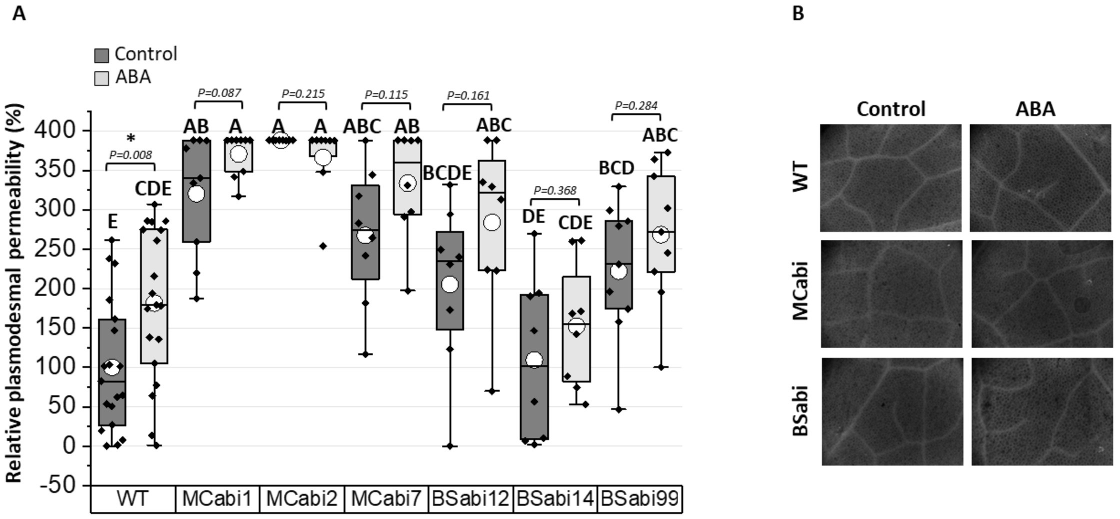


**Supplementary Figure 8.** The symplastic pathway's response to ABA. (A) DANS assays of the WT and all independent MCabi and BSabi lines petiole-fed AXS (dark gray) or AXS + 10 μM ABA (light gray) showing cell-to-cell movement of the fluorescent dye. (B) A representative picture of callose staining of the same leaves; callose was the positive control for the fluorescent staining.

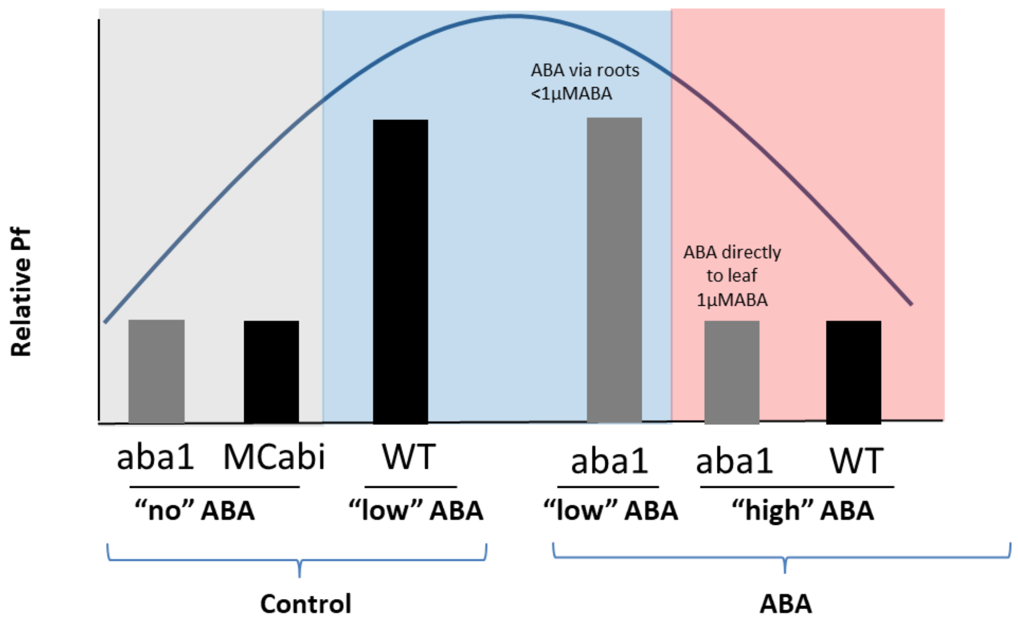


**Supplementary Figure 9.** Proposed dose-response of spongy MC $P_f$ to ABA. This figure shows summarized data from Morillon and Chrispeels (2001), which addressed the mesophyll in general, without distinguishing between spongy and palisade mesophyll (*aba1*, gray bars), as well as data from our current study, which addressed the spongy mesophyll (MCabi#2, black bars). When ABA signaling/sensing is low, due to loss of synthesis (*aba1*) or signaling (MCabi), $P_f$ is at its lowest levels (gray area). In the presence of low levels of ABA (e.g., WT basal levels of ABA), spongy MC $P_f$ is relatively high. Similar high $P_f$ was observed for the ABA-nonproductive *aba1* plants when 1 µM ABA was supplied via the roots (presumably resulting in low levels of ABA in the leaf blue area). However, higher levels (1 µM ABA applied directly to WT protoplasts or to the leaves of *aba1*) resulted in low *Pf* levels (pink area).

**Supplemental Table 1**. Results of the two-way ANOVA.

| | | | E | | | Ψw | | | $K_{leaf}$ | | | $g_s$ | | | $A_N$ | | | iWUE | | |
|---|---|---|---|---|---|---|---|---|---|---|---|---|---|---|---|---|---|---|---|---|
| **Fig. 2** | Source | DF | Mean Square | F Ratio | Prob > F | Mean Square | F Ratio | Prob > F | Mean Square | F Ratio | Prob > F | Mean Square | F Ratio | Prob > F | Mean Square | F Ratio | Prob > F | Mean Square | F Ratio | Prob > F |
| | Treatment | 1 | 88.2545 | 208.637 | 0001.> | 0.0659587 | 4.4636 | 0.0365 | 6358.542 | 40.1832 | 0001.> | 1631544 | 216.243 | 0001.> | 138.7754 | 152.328 | 0001.> | 234.9315 | 92.5466 | 0001.> |
| | Line | 2 | 23.30837 | 55.102 | 0001.> | 0.0600436 | 4.0633 | 0.0194 | 2285.8 | 14.4452 | 0001.> | 377818 | 50.0757 | 0001.> | 24.0971 | 26.4506 | 0001.> | 93.3172 | 36.7605 | 0001.> |
| | Line*Tre | 2 | 1.57261 | 3.7177 | 0.0268 | 0.0097369 | 0.6589 | 0.5191 | 400.441 | 2.5306 | 0.0835 | 79501 | 10.537 | 0001.> | 2.4861 | 2.7289 | 0.0688 | 9.4095 | 3.7067 | 0.027 |
| | Error | 138-132 | 0.423 | | | 0.014777 | | | 158.24 | | | 7545 | | | 0.911 | | | 2.5385 | | |
| | C. Total | 143-137 | | | 0001.> | | | 0.0236 | | | 0001.> | | | 0001.> | | | 0001.> | | | 0001.> |

| | | | Relative PD permeability | | |
|---|---|---|---|---|---|
| **Fig. 6** | Source | DF | Mean Square | F Ratio | Prob > F |
| | Treatment | 1 | 108178.5 | 13.4396 | 0.0004 |
| | Line | 2 | 467873.1 | 58.1263 | 0001.> |
| | Line*Tre | 2 | 6526.1 | 0.8108 | 0.4467 |
| | Error | 132 | 8049 | | |
| | C. Total | 137 | | | 0001.> |

| | | | MCabi E | | | MCabi Ψw | | | MCabi Kleaf | | | | BSabi E | | | BSabi Ψw | | | BSabi Kleaf | | |
|---|---|---|---|---|---|---|---|---|---|---|---|---|---|---|---|---|---|---|---|---|---|
| **Supp. Fig. 2** | Source | DF | Mean Square | F Ratio | Prob > F | Mean Square | F Ratio | Prob > F | Mean Square | F Ratio | Prob > F | DF | Mean Square | F Ratio | Prob > F | Mean Square | F Ratio | Prob > F | Mean Square | F Ratio | Prob > F |
| | Treatment | 1 | 102.2933 | 103.742 | 0001.> | 0.0803454 | 4.3715 | 0.038 | 9912.337 | 21.1477 | 0001.> | 1 | 64.4526 | 88.5485 | 0001.> | 0.04688 | 4.4437 | 0.0374 | 2728.514 | 33.2814 | 0001.> |
| | Line | 3 | 25.3971 | 25.7568 | 0001.> | 0.017307 | 0.9416 | 0.4218 | 1968.056 | 4.1988 | 0.0067 | 3 | 5.3061 | 7.2898 | 0.0002 | 0.03263 | 3.0922 | 0.0303 | 434.718 | 5.3025 | 0.0019 |
| | Line*Tre | 3 | 0.4188 | 0.4247 | 0.7355 | 0.0124275 | 0.6762 | 0.5677 | 109.914 | 0.2345 | 0.8722 | 3 | 1.27499 | 1.7516 | 0.161 | 0.00170 | 0.1613 | 0.9222 | 128.024 | 1.5616 | 0.2032 |
| | Error | 181-175 | 0.986 | | | 0.018379 | | | 468.72 | | | 105-104 | 0.7279 | | | 0.01055 | | | 81.983 | | |
| | C. Total | 188-182 | | | 0001.> | | | 0.2139 | | | 0001.> | 112-111 | | | 0001.> | | | 0.0548 | | | 0001.> |

| | | | Relative PD permeability | | |
|---|---|---|---|---|---|
| **Supp. Fig. 8** | Source | DF | Mean Square | F Ratio | Prob > F |
| | Treatment | 1 | 76404.4 | 11.1141 | 0.0011 |
| | Line | 6 | 187470.2 | 27.2702 | 0001.> |
| | Line*Tre | 6 | 5672.5 | 0.8251 | 0.5525 |
| | Error | 124 | 6874.5 | | |
| | C. Total | 137 | | | 0001.> |

### MATERIALS AND METHODS

*Quantification of abi1-1 Expression by Restriction*

*abi1-1* expression was quantified by restriction, as described by Negin et al. (2019). In brief, total RNA was extracted from leaves and cDNA was prepared. The ABI1_174F (TCTGGGTCACATGGTTCTGA) and ABI1_1064R (CCATCTCACACGCTTCTTCA) primers were used for amplification of both ABI1 and *abi1-1*. The amplified 911-bp fragment was cleaned and 1000 ng of the amplified fragment were digested for 2 h by NcoI and loaded onto a 1% agarose gel. The gel was photographed under UV light and digital image analysis was performed using ImageJ software (http://rsb.info.nih.gov/ij/).

*Foliar ABA Concentrations*

Concentrations of ABA in the leaves were measured as described by Negin et al. (2019). Briefly, leaves were immediately weighed (150–400 mg) and transferred to -80°C. Foliar ABA was extracted, purified and quantified by a liquid chromatography-mass spectrometry system that consisted of a Dionex Ultimate 3000 RS HPLC coupled with a Q Exactive Plus Hybrid Fourier Transform mass spectrometer equipped with a heated electrospray ionization source, using an internal standard.
